# Archaic humans have contributed to large-scale variation in modern human T cell receptor genes

**DOI:** 10.1101/2022.08.25.505097

**Authors:** Martin Corcoran, Mark Chernyshev, Marco Mandolesi, Sanjana Narang, Mateusz Kaduk, Christopher Sundling, Anna Färnert, Carolina Bernhardsson, Maximilian Larena, Mattias Jakobsson, Gunilla B. Karlsson Hedestam

## Abstract

The human T cell receptor (TCR) genes are critical for mediating immune responses to pathogens, tumors and regulating self-antigen recognition. A detailed analysis and validation of expressed TCR alpha, beta, gamma, and delta genes in 45 donors from 4 human populations: African, East Asian, South Asian, and European, revealed a total of 175 novel TCR variable and junctional alleles. The majority of novel alleles contained coding changes and were present at widely differing frequencies in the populations, a finding confirmed using DNA samples and sequences from the 1000 Genomes Project. Importantly, we identified three Neanderthal-derived, introgressed TCR regions, including a highly divergent novel TRGV4 variant, present in all archaic assemblies, that was frequent in all modern Eurasian population groups. Our results demonstrate significant variation in TCR genes at both individual and population levels, providing a strong incentive for including allelic variation in studies of TCR function in human biology.

## Introduction

Genetic diversity in components of immune response pathways is increasingly seen as the normative state in outbred species (1–4). The extent of this diversity and the likely biological implications of specific immunological variations is, to date, unclear. Host-pathogen interactions may result in negative, positive, or balancing selection within components of the immune system (5), resulting in allelic diversity at the level of populations. While variation within the innate immune system had been extensively studied (4), the adaptive immune system remains largely unexplored at the level of individual and population-based variation, with many human population groups severely under-represented in reference databases of immune genetic variants (6).

The adaptive immune system of all jawed vertebrates can be considered to function as a tripartite system. In addition to B cells, which can differentiate into antibody-producing cells, it consists of CD8+ or CD4+ alpha/beta (αβ) T cells that recognize foreign peptides presented by major histocompatibility complex (MHC) class I or class II molecules, respectively. The third group of lymphocytes are gamma/delta (γδ) T cells that function independently of MHC presentation and are known to recognize lipids, phosphoantigens and intermediates of the isoprenoid biosynthesis pathway (7, 8). Following selection in the thymus, γδ T cells play diverse roles in the surveillance of infections and cellular stress (9). A primary modulator of αβ T cell function is the MHC background of the individual, with over 16,000 class I and 6,000 class II allelic variants known to date (10). While TCR antigen recognition by αβ T cells involves close interactions with the highly polymorphic MHC class I and class II molecules, the roles and extent of TCR germline gene polymorphisms for shaping these interactions remain unknown.

Like the immunoglobulin loci, the TCR loci consist of arrays of variable (V), diversity (D) and junctional (J) genes that recombine during T cell development to produce a unique functional TCR with specific binding potential. Both the immunoglobulin and TCR VDJ genes have expanded through genomic duplication events over evolutionary time with the result that the current loci contain, in addition to the functional genes, numerous non-functional pseudogenes or open reading frames (ORFs) that lack functional recombination sequences (11, 12). The sequence of functional and pseudo-/ORF genes are highly similar over significant nucleotide stretches, which creates difficulties for short read sequence assembly and subsequent identification of variant alleles (13). This factor has inhibited the identification of TCR gene variation as most genome analyses have utilized short read sequencing technologies and TCR repertoire sequencing (Rep-seq) projects have concentrated on incomplete TCR sequences, with the identification of complementarity determining regions 3 (CDR3s) being the primary goal of such studies (14). In addition to this, the current knowledge of TCR germline gene variation is limited to those identified in a limited set of studies and genomic assemblies. Most of the current the international ImMunoGeneTics information system^®^ (IMGT) TCR database is derived from just two genomic assemblies, GRch37/hg19 and GRch38/hg38, that are primarily European in terms of allelic content.

Here, we utilized an alternative approach to delineate TCR gene variation in 45 individuals from multiple human populations. Using Rep-Seq analysis and germline allele inference with the program IgDiscover (15), all known and novel TCR alpha (TRA), beta (TRB), gamma (TRG) and delta (TRD) allelic variants from each of 45 individuals were identified. We also undertook an extensive series of validations to ensure accurate TCR genotyping in each of the 45 individuals. We further analyzed publicly accessible TCR Rep-seq libraries from five monozygotic twin pairs and showed that many of the novel alleles were present in this independent dataset. The collection of the full set of expressed alleles from a diverse population set enabled the assembly of a human TCR VDJ germline gene database, identifying alleles that varied in frequency between different populations. Critically, our studies enabled the comparison of the full set of alleles with those identified in archaic human high coverage assemblies, revealing several introgressed genomic segments that are present in in modern humans.

## Results

A total of 45 individuals who self-identified as members of either sub-Saharan African, East Asian, South Asian, or European populations provided blood samples that were used to isolate peripheral blood mononuclear cells (PBMCs). Total RNA isolated from these cells was used to create TRA, TRB, TRG and TRD libraries using a 3’ constant region-located unique molecular identifiers (UMIs) and a 5’ multiplex library amplification procedure. Each of the 5’ primers targeted the leader regions of the appropriate genes, thereby enabling full-length V(D)J amplification, a prerequisite for high confidence V and J germline gene inference using the IgDiscover software (15) (**Figure 1A**). The analysis pipeline resulted in the identification of known and novel V and J alleles from each of the four loci and the production of a complete TRA, TRB, TRG and TRD genotype for all 45 individuals using the IgDiscover *corecount* module. These analyses enabled the production of a combined allelic database of expressed V and J alleles from the 45 donors for all four TCR loci (**Figure 1B, SI Figure 1A-C**). A total of 134 expressed TRAV alleles, were identified across the cases analyzed, of which 76 (56.7%) were novel (**Figure 1B**). A single novel TRDV-exclusive allele, TRDV2*02_S6129, was identified, and is an extension of TRDV2*02, an allele that contains a 3’ truncation in the IMGT reference database. Eight novel alleles, however, were found in 3 different V genes shared by both the TRA and the overlapping TRD locus (**Figure 1B**). In addition, 131 TRBV alleles and 27 TRGV alleles were identified, of which 65 (49.6%) and 15 (55.5%) were novel, respectively (**Figure 1B**). Novel expressed alleles, not present in the IMGT database, were identified for 30/43 TRAV genes, 33/48 TRBV genes and 7/8 TRGV genes (**Figure 1C**). Of note, 22 novel alleles were found that encompassed sequences present in the IMGT database, indicating erroneous truncation of between 2 to 35 nucleotides (nts) for those alleles found within the IMGT database.

**Figure 1.**
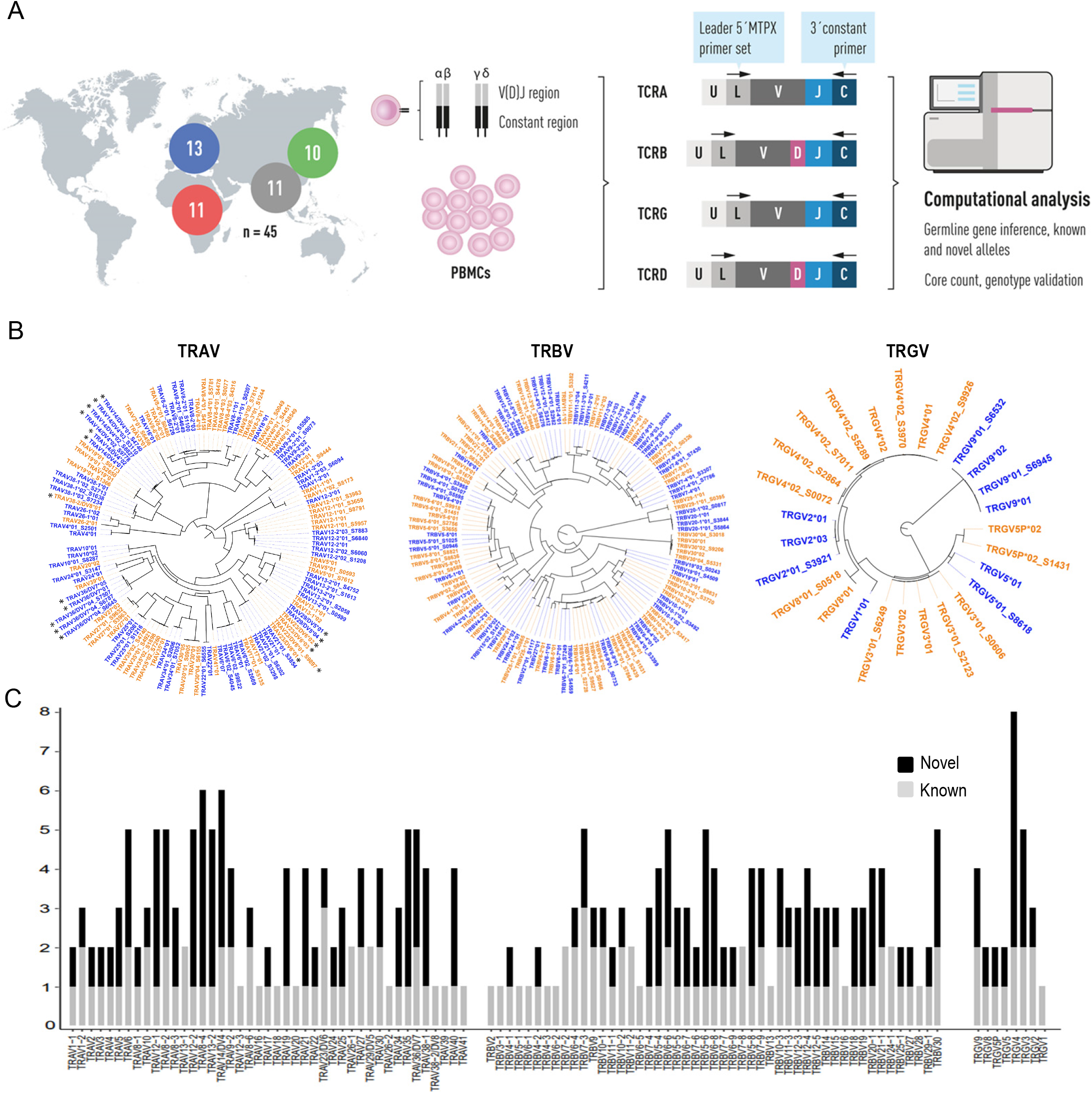
**A.** Procedure for genotypic analysis of expressed TCR genes in volunteers from multiple human population groups. Total RNA isolated from 45 individuals was used in the preparation of TRA, TRB, TRD and TRG NGS libraries for each case. 11 samples derived from individuals from Africa, 13 from Europe, 11 from South Asia and 10 from East Asia. The libraries utilized a 3’ UMI incorporated during cDNA synthesis using a constant region primer and a 5’ multiplex set of forward primers located within the V gene leader sequence that enabled full V(D)J amplification of all expressed genes. The libraries were sequenced using an Illumina MiSeq instrument and the germline identification performed using the program IgDiscover. Additional genotype analysis was performed with the *corecount* module of IgDiscover. **B.** Dendrograms of the combined allelic databases identified for the entire set of TRAV, TRBV and TRGV alleles expressed in the 45 cases. In all dendrograms the color alternates between red and blue for easier identification of different genes. Expressed alleles identical to known TCR V alleles are denoted with standard IUIS TCR allelic nomenclature. Novel alleles are denoted by the presence of a four numbered suffix. Novel TRDV alleles found in genes that are jointly utilized by TRA and TRD are denoted with an asterisk. **C.** Stacked bar chart illustration of the numbers of known and novel TRAV, TRBV and TRGV allelic variants identified per gene in the 45 cases. Known alleles are shown in grey and novel alleles shown in black. The x-axis represents TRAV, TRBV and TRGV genes in the chromosomal order

TCR V alleles with coding changes comprised the majority (97/153, 63.4%) of novel V alleles identified (**SI Figure 1D**). The percentage of coding changes in novel J alleles was even more striking, with 20/20 (100%) of the novel TRAJ and TRBJ alleles resulting in an amino acid change in the translated sequence compared to the known alleles of the same gene (**SI Figure 1D-F**). The frequency of allelic variation of both V and J genes enabled inferred haplotype analysis to be performed in many cases, revealing novel alleles to be present in both heterozygous and homozygous form (**SI Figure 2B**).

### Validation of inference process

While germline inference from expressed B cell receptor (BCR) Rep-seq data is an established approach to identify novel immunoglobulin (Ig) alleles (15–18), adaptive immune receptor repertoire (AIRR) sequencing combined with germline inference was to date only reported in a recent study that focused on TRB allelic variation in one population group (19). The IgDiscover inference process is based on the identification of germline V, D or J sequences through their use in multiple independent V(D)J recombination events, and the exclusion of erroneous sequences, such as those caused by PCR error. To this end we modified the original IgDiscover tool to account for TCR specific features and undertook a series of independent validation approaches to ensure accuracy of the novel alleles and individualized genotypes presented here.

We used Sanger sequencing to validate an extensive set of both previously known and novel alleles identified from the expressed TCR libraries using targeted PCR across the specific gene loci. A total of 32 novel alleles were validated from the donor set, of which five were novel TRAJ, 15 novel TRAV, one novel TRBJ, eight novel TRBV and three novel TRGV alleles. To show that alleles identified in the donor cases were found in the wider population we identified sets of single nucleotide polymorphism (SNP) variants specific for each novel allele. We then identified individual cases from the 1000 Genome Project (1KGP) that potentially contained these alleles and used the corresponding DNA as template for PCR and Sanger validation of 52 novel alleles identified in the 45 donors. Of these six were novel TRAJ, 27 novel TRAV, 16 novel TRBV and three novel TRGV alleles. Of these, 11 novel alleles were Sanger validated in both donor and 1KGP samples (**SI Figure 1A**).

IgDiscover analysis of two familial groups, A and B, from our donor set enabled the use of familial inheritance as a means of obtaining additional evidence that the inference process was accurate at both the genotypic and haplotype level (**SI Figure 2A**). Familial group A comprised two parents (D25, D45) and one child (D05), while group B comprised a separate parent and child (D06 and D08 respectively). For familial group A, nine novel TRAV alleles were inherited by the child (**SI Figure 2B**) in addition to seven novel TRBV alleles. In the case of familial group B, eight novel alleles are shared between the parent and child, five TRAV and three TRBV. We also utilized a previously published independent dataset of TRA and TRB Next Generation Sequencing (NGS) libraries from five sets of monozygotic twins (20). We were able to identify five novel TRAJ, 18 novel TRAV and 10 novel TRBV alleles present in both members of one or more of the five sets of twins. Of these, one TRAJ allele (TRAJ27*01_S4866), four TRAV alleles (TRAV13-1*02_S8036, TRAV21*01_S8011, TRAV38-1*01_S2525 and TRAV4*01_S2986), and one novel TRBV allele (TRBV5-4*01_S3690) were found only in the twin cases.

Inferred haplotype analysis of expressed sequences provides strong support for V, D and J germline inferences (21, 22) and we used this approach extensively in this study. A total of 95 novel alleles had haplotype validation, of which 47 novel TRAV alleles were heterozygous and 18 were homozygous. Sixteen novel TRAJ alleles were heterozygous in the 45 donor cases and an additional allele, TRAJ27*01_S4866, was found in twin pair 2. A total of 24 novel TRBV alleles were found to be heterozygous in donor cases and three homozygous alleles, TRBV5-5*01_S1025 in D09 and D43 and TRBV18*01_S1676 in D29 and TRBV20-1_S5864 in case D15. In total, three novel TRBJ alleles were found to be heterozygous by haplotype analysis. The TRG locus has a lack of suitable heterozygous alleles that can function as haplotyping anchors and so only one novel V allele, TRGV2*01_S3921, was haplotyped as heterozygous, as well as a single novel J allele, TRGJP2*01_S8334, in both D28 and D32.

The identification of the same, full-length novel allele in multiple individuals provides additional strong supportive evidence that the sequence is germline-encoded. This approach, applied to the 45 donor cases and five sets of monozygotic twins, revealed a total of 42 novel TRAV alleles found in more than one individual, eight novel TRAJ alleles, 38 novel TRBV alleles, and seven novel TRGV alleles. In addition, one novel TRGJ allele, TRGJP2*01_S8334, was found in more than one individual. In many cases the novel TCR alleles were found in multiple individuals. TRAV8-4*03_S4316 was found in 23 individuals. As mentioned above, several frequently inferred novel alleles encompass known alleles, such as TRAV35*01_S4921 and TRAV36/DV7*04_S7507, which are longer versions of TRAV35*02 and TRAV36/D7*04. This reveals that those two known alleles are erroneously truncated in the current IMGT reference sequence database. A comparative analysis of the current IMGT database with the alleles identified here resulted in a set of 25 alleles in which the truncation was corrected through the current study.

The function of a given TCR may be affected by any variation in germline alleles that produces amino acid changes since TCR genes unlike BCR genes do not undergo somatic hypermutation. The coding changes seen in 84/136 novel alleles occurred in both the CDRs and framework regions (FR), including the hypervariable region 4 (HV4). Even more striking was the coding variation found in the TCR J genes, with 21/24 (87.5%) of novel J alleles showing coding variation compared to the J alleles in the current IMGT database. All 17 novel TRAJ and four novel TRBJ alleles were found to have coding variation, with 6/17 and 2/4 having the coding differences within the CDR3 region of the TRAJ and TRBJ novel alleles, respectively (**SI Figure 1D-F**).

### Loss of expression is frequently associated with loss-of-function allelic variants

High resolution characterization of individualized sets of expressed VDJ germline alleles, and the ability to perform haplotype analysis on many of the cases studied here, enabled TCR genotype comparison at the gene and allelic level. This analysis revealed a higher level of heterozygosity in the TRAV locus compared to the TRBV locus (**Figure 2A-B**). Gene duplications were rare, but hemizygous or homozygous loss of expression was apparent for several genes. The previously described frequent 21.5 kb genomic deletion encompassing TRBV4-3 (23) was visible in the expressed genotypes, with 16 cases showing homozygous loss and three cases showing hemizygous loss. For several genes, loss of expression was found to be associated with genetic variants that are expected to inhibit function. For example, TRAV26-2 shows frequent hemizygous (six cases) and homozygous (five cases) expression loss (**Figure 2A**). We found that a SNP variant, rs2242542 G/A, located at position 2 of the splice acceptor AG sequence at the 5’ end of the TRAV26-2 V exon, where the ‘A’ variant removes the splice site, was present at a similarly high frequency of 29.1% in the 1KGP cases (1458/5008). Furthermore, sequence variants that result in stop codons, rs54770561, rs17113205, rs768942453, rs17249 and rs361451, were identified in five V genes that show hemizygous expression loss, namely TRAV1-2, TRAV19, TRAV34, TRBV10-1 and TRBV6-8 respectively (**Figure 2**). Three alleles containing stop codons were Sanger validated, while low level expression of stop codon-containing alleles was found for all five V genes in libraries that appeared as deletion cases. We designated alleles containing stop codons, but which were expressed at the RNA level at low but detectable levels within the libraries, expressed non-functional (ENF) alleles. Variation was limited in TRDV and TRDJ in contrast to TRGV (**SI Figure 3A-B**). In the latter, one individual, D12, showed loss of expression of both TRGV4 and TRGV5 (**SI Figure 3B**). A multiplex PCR analysis using primers specific for TRGV4 and the surrounding genes, TRGV2 and TRGV9, confirmed homozygous genomic deletion of TRGV4 in this case (**SI Figure 3C**).

**Figure 2.**
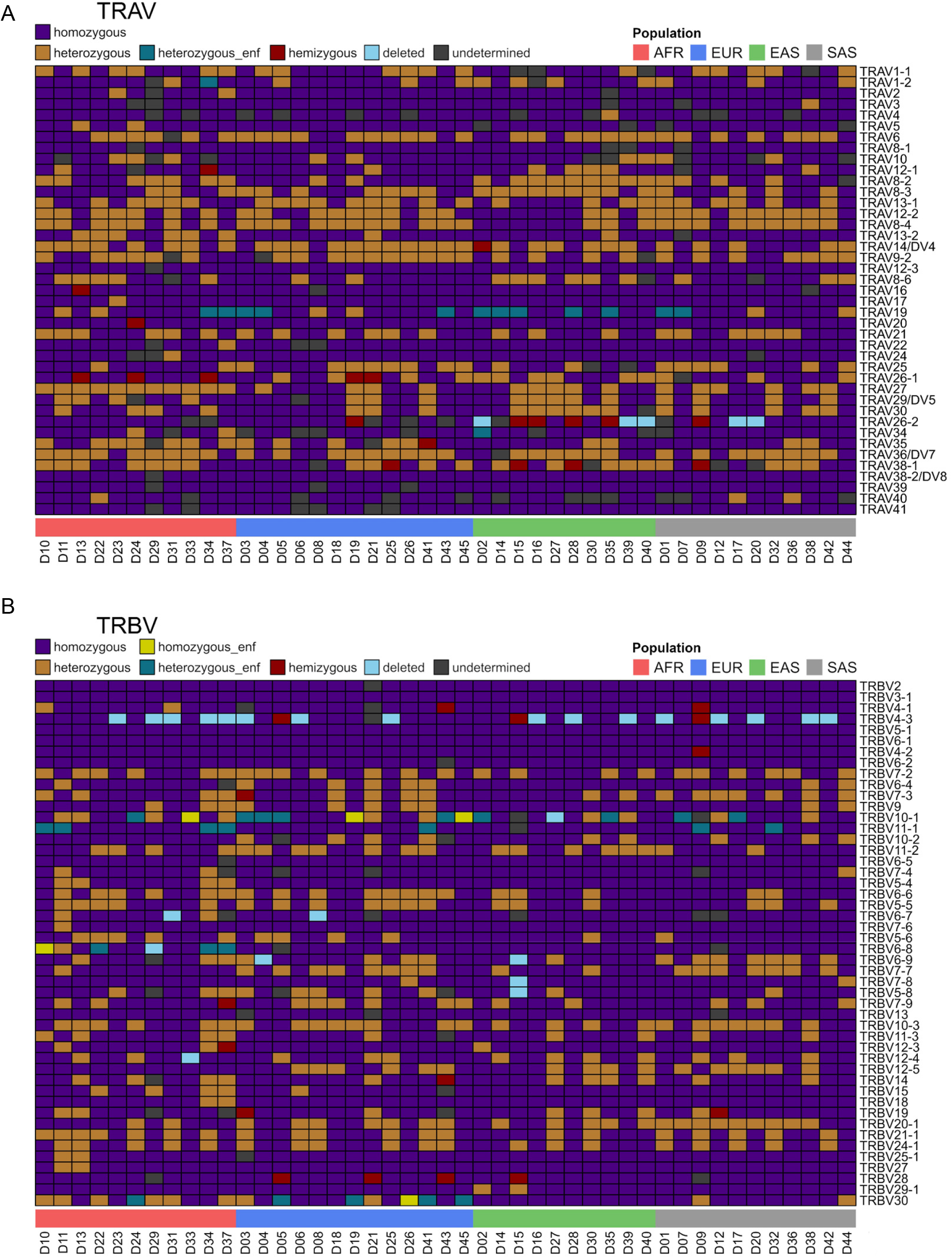
**A**. Illustration of TRAV and **B**. TRBV variation at the gene level. Inferred haplotype analysis identified the presence of homozygosity, heterozygosity, hemizygous, homozygous, or expressed non-functional (ENF) presence that are denoted by blocks of color, the key being present on the top left above the figure. The population groups of the case are shown below the figure, the key being present on the top right of the figure. For each case the gene names are on the right-hand side and the case numbers are shown below the figure. The genes are represented on the y-axis in chromosomal order.

Loss of expression of three TRAJ genes was also found to be associated with SNP variants that affect recombination, splicing or disruption of the reading frame of the genes. The TRAJ12 gene was lost in homozygous form in two cases, (D13 and D31) and in hemizygous form in two cases (D10 and D34) (**SI Figure 4A (upper panel)**). A common SNP variant, G/A rs62622786, was found to be located within the recombination signal sequence (RSS) of this gene (**SI Figure 4B (upper panel)**). An additional deletion variant, rs3841038 AAGT/− removes the splice donor site of TRAJ17, a gene that showed hemizygous (two cases) and homozygous (one case) loss of expression, respectively (**SI Figure 4A (lower panel), SI Figure 4B (lower panel)**). Hemizygous loss of expression of TRAJ52 was found in two cases, D02 and D27, both of which showed low level expression of a 1 nt shortened TRAJ52 sequence that corresponds to the A/− SNP variant rs143418056. In the TRA libraries from D02 and D27, the SNP-deleted TRAJ52*01_S5131 allele could be haplotyped using heterozygous TRAV26-1 and TRAV14/DV4 alleles respectively as anchors. This analysis revealed that the TRAJ52*01 allele was combined with different heterozygous TRAV26-1 and TRAV14/DV4 alleles respectively, to those recombined with the SNP-deleted TRAJ52 allele (**SI Figure 4C**).

### Allelic and structural variation in population groups

Standardized library preparation for the 45 donor cases in this study enabled the comparison of individualized genotypes. The resulting genotypic data revealed significant allelic variation at an individual level. Allelic heterozygosity is common in the TRA, TRB and TRG loci (**Figure 3A-E**). Most alleles found in the TRA, TRB, TRG and TRD loci are present in all the four population groups analyzed (**Figure 4A**). However, in each locus we found variation in allele frequencies, where certain population groups carried some exclusive alleles. We identified 30, seven, seven and four TRAV alleles solely in the African, East Asian, South Asian, and European groups, respectively. For TRBV we found 33, four, two and six TRBV alleles solely in the African, East Asian, South Asian, and European groups, respectively. For TRGV, we found eight, one, two, and one TRGV alleles solely in the African, East Asian, South Asian, and European groups, respectively. For the TRAJ genes we identified five, three, and one TRAJ alleles solely in the African, East Asian, and European groups, respectively. Finally, for TRBJ we identified three and one TRBJ alleles solely in East Asian and South Asian cases, respectively. These observations are consistent with generally greater genomic diversity among sub-Saharan Africans compared to non-African populations (24, 25).

**Figure 3.**
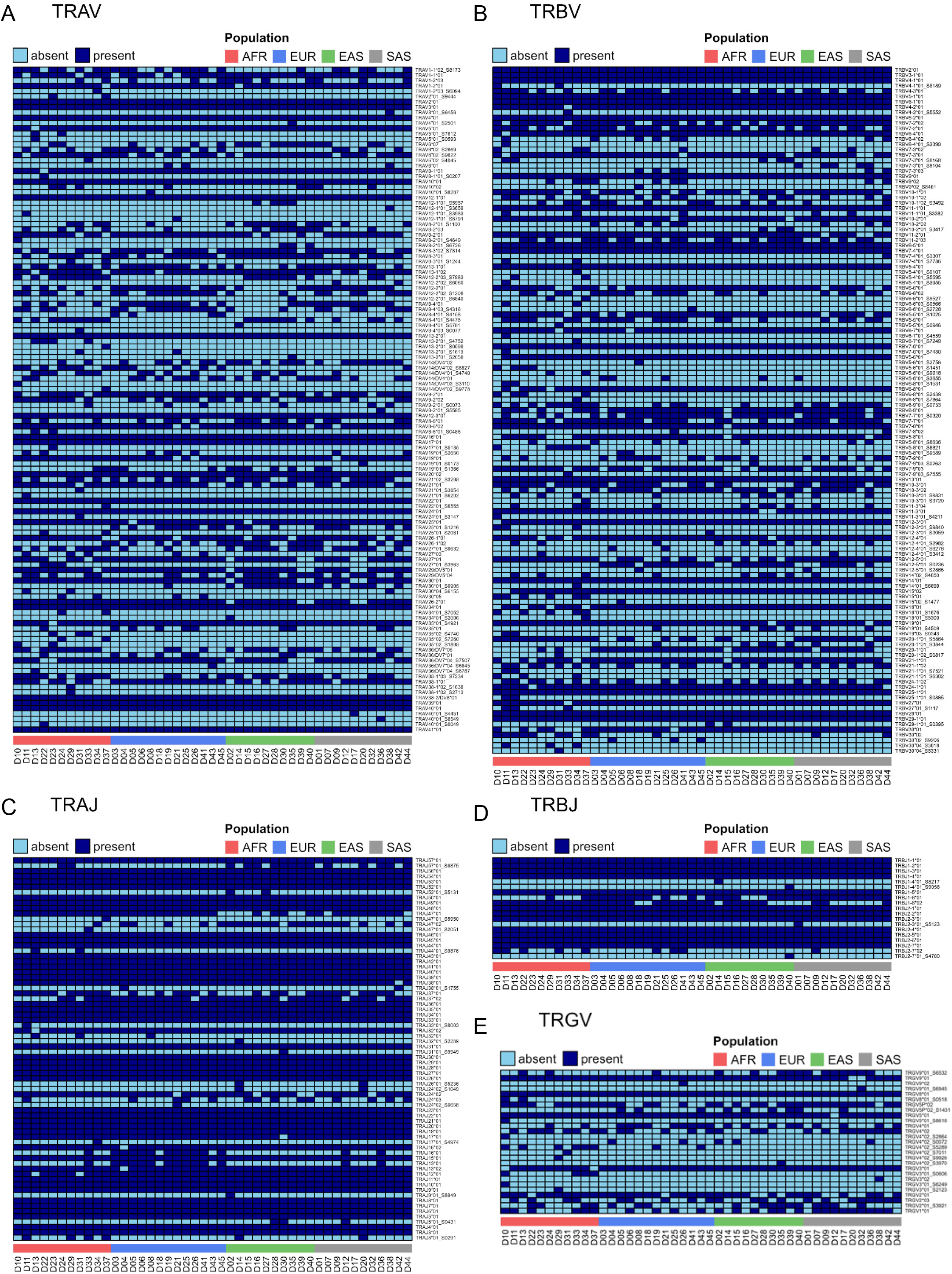
**A**. TRAV allelic variation **B**. TRBV allelic variation **C**. TRAJ allelic variation **D**. TRBJ allelic variation **E**. TRGV allelic variation. For all the panels, the presence (dark blue box) or absence (light blue box) of each allele in the 45 donors are indicated. The allele names are present on the right side of each figure and donors are indicated in columns and grouped according to indicated populations.

**Figure 4.**
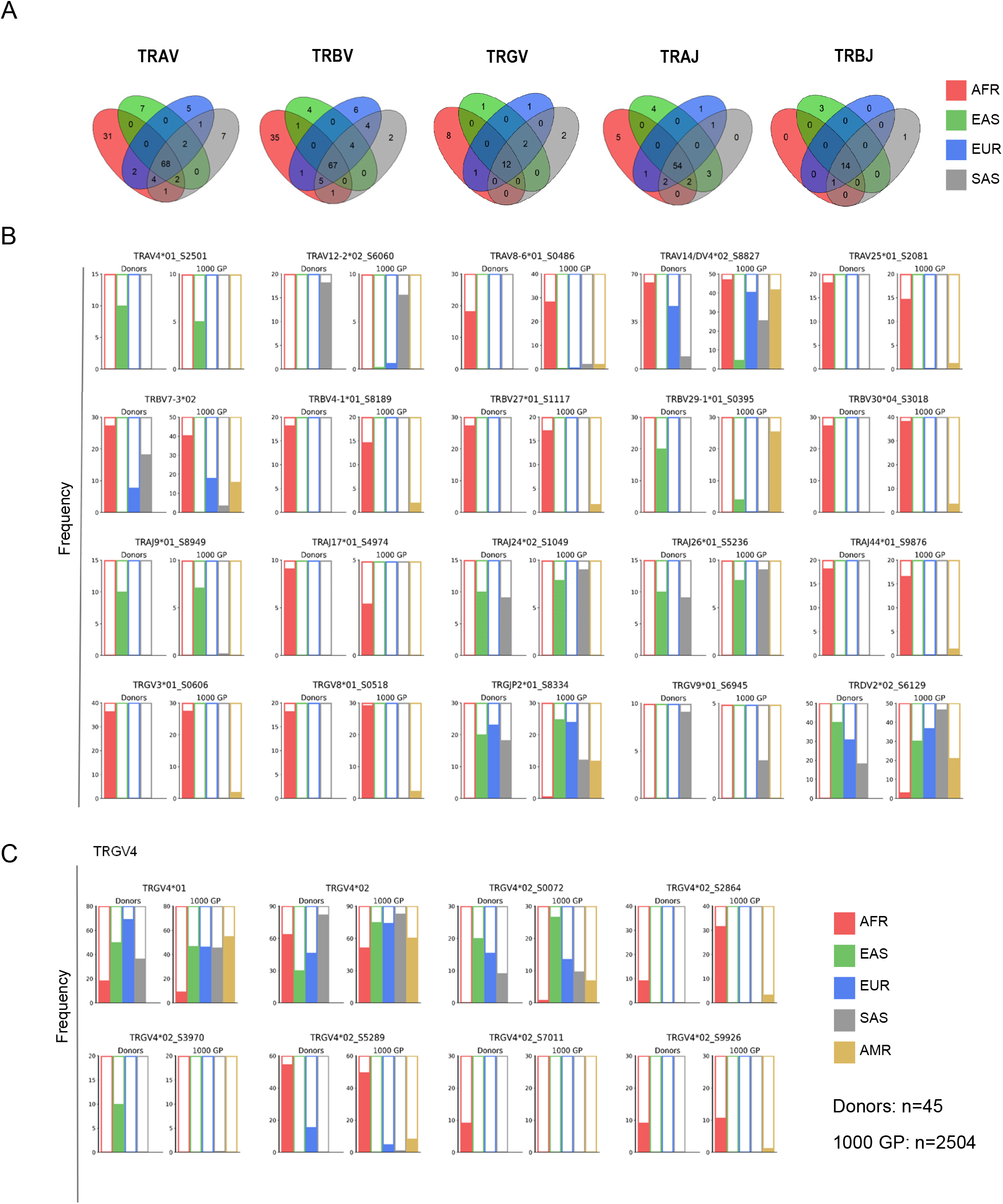
**A.** Venn diagram of numbers of individual TCR alleles found in the set of individuals from African, East Asian, South Asian, and European population groups. From left to right, TRAV, TRBV, TRGV, TRAJ, and TRBJ genes. The population key is shown on the right of the figure. **B.** Variation in the frequency of TCR alleles in different human populations. For each allele the left bar chart shows the frequency of the allele in the donors from populations analyzed in this study and the right bar chart shows the frequency of allele specific SNPs within the set of 2504 samples from the 1000 genomes set. **C**. Variation in the frequency of TRGV4 alleles in the analyzed population samples and in the frequency of allele associated SNPs in the full set of the 1000 genome samples.

Many of these alleles, however, were only found in single individuals in our analysis, and may be rare within the population at large. To strengthen these results, we identified SNP variants specific for each allele found in our analysis and used these to determine the likely presence of the allelic variants within the full 2504 individuals of the 1KGP dataset (http://www.ensembl.org/info/website/tutorials/gene_snps.html). Once such allele-specific cases were identified, we confirmed the presence of a selected group of alleles using the appropriate 1KGP DNA sample for targeted genomic PCR followed by cloning and Sanger sequencing. The SNP combination results in the 1KGP cases show a strong agreement between the population frequency of the alleles identified in our 45 donors. This analysis provided an additional use as it allowed for the determination of the allelic frequency in an additional continental group, the American population, which was absent from our donor set.

We identified 32 alleles present in more than one individual from the African population group in our study, which were absent from individuals of all other populations (**Figure 4A**). This includes one TRAJ, 11 TRAV, 17 TRBV and two TRGV alleles. One TRAV allele and three TRBV alleles were found in more than one European individual and not in individuals from other groups. For the East Asian group, we identified two TRAV and two TRBV alleles in more than one individual that were not found in other population groups. Finally, for the South Asian group, we found one TRAV and one TRBV allele in more than one individual, which were absent from all other population groups. We additionally identified several alleles that were present in two populations but missing from the others.

In addition to alleles present in specific populations, we identified several common alleles that were absent in a particular population. Despite the African group showing the highest number of alleles overall, we identified three TRAV, one TRBV, one TRDV, one TRGJ and two TRGV alleles that were found in other populations, but which were absent among the African individuals. Likewise, we identified one TRAJ and one TRAV alleles that are found in other populations but absent from the European cases (**Figure 4A**). For some of the novel alleles described in this study, the difference in frequencies between the population groups was striking. TRAJ44*01_S9876 was found in 2 of 11 African individuals (cases D13 and D31) and rs76402200 T, the SNP variant specific for this allele, was found at a similar percentage of 1KGP cases from the same population (**Figure 4B**). TRBV30*04_S3018 was found in almost 30% of African individuals and the allele specific SNP combination, rs17269 G, rs361463 A, rs361462 T and rs361461 T, was found in almost 40% of African 1KGP cases (**Figure 4B**). TRGV3*01_S0606 and TRGV8*01_S0518 were likewise found in almost 40% and 20% of African donors, respectively (**Figure 4B**).

For several genes we found multiple alleles that appeared to be population specific. For example, the African specific variants, TRGV4*02_S2864, TRGV4*02_S5289 and TRGV4*02_S9926, were present for the same gene that contains the variant TRGV4*02_S0072, which is almost absent from the African group but is relatively frequent in the European, South Asian, and East Asian groups (**Figure 4B**). Additionally, TRBV19*01_S4509 was found in East Asian, South Asian, and European individuals while TRBV19*03_S0243 was found only in African individuals.

The large numbers of novel TCR alleles found in non European populations supports the idea that immunological genetic variation amongst many populations has not been sufficiently documented (6). Indeed we obtained additional evidence of the high levels of TCR variation by utilizing genomic sequence of human BAC clones to cross validate many of the African associated novel alleles. For example, BAC clones RP11-114B6 and RP155E15 contain four and eight novel TRBV alleles, respectively, that were also found in African donors in our study. These novel alleles comprised 50% (4/8) and 44% (8/18) of the total number of functional TRBV genes on these BAC genomic clones. An additional 5 novel TRBV alleles and two novel TRGV alleles identified in our donors are also found in a different set of three BAC clones. (**SI Figure 5A-B**). It is also striking that loss of function alleles underly much of the population specific variation. TRAJ12 and TRAJ17, in African and East Asian cases, respectively, contain loss of function variants as discussed above that lead to hemizygous or homozygous loss of expression (**SI Figure 4A-B)**. TRAJ52*01_S5131 in East Asian cases D02 and D27 also contains a single nucleotide deletion that results in a frameshift and stop codon that greatly reduced expression and was present in heterozygous form in two cases (**SI Figure 4C**). Likewise, we found ENF alleles underlying loss of expression of TRBV6-8 and TRBV30 in African and European cases, respectively.

No novel large common structural alterations were identified in the 45 cases analyzed. In case D15, a possible genomic deletion was found in the TRBV locus, as three contiguous genes, TRBV5-8, TRBV7-8, and TRBV6-9 were absent in the expression data of this donor **(Figure 2).** The previously described TRBV4-3 deletion was found in 19/43 cases (**Figure 2**), a genomic deletion also associated with concomitant loss of TRAV7-2*02 in 13/16 cases (**Figure 3A**). However, in most instances of apparent gene loss, the absence of expression was associated with the presence of ENF alleles that contain stop codons.

### Archaic introgressed regions within the TCR loci

The immune system of *Homo sapiens* has evolved within the context of the specific environments they inhabited (26). This environment has been almost exclusively within sub-Saharan Africa until approximately 80,000 years ago when some modern humans migrated out of Africa into Eurasia and subsequently into the far reaches of Australia, the Americas, and the Pacific Islands. Strong evidence suggests that along the course of these migrations these ancestors of modern humans encountered and interbred with members of their evolutionary cousins, the Neanderthals and the Denisovans, whose ancestors had migrated from Africa more than 500,000 years previously (27). Variants of adaptive immune receptor genes specific to these archaic populations may have entered the modern human gene pool during such encounters. The production of an extended database of TCR VDJ alleles that could be linked with specific human populations enabled us to ask whether any of the variants we identified could have been introduced through introgression events. To investigate this, we performed a comparison of the VCF-derived TCR gene sequences from four high coverage archaic reference assemblies of three Neanderthals, namely Vindija (28), Chagarskya (29) and Altai (30), and the single Altai Denisovan (31).

The results revealed that many alleles are shared between modern and archaic humans, indicating a long evolutionary history of much of the current human TCR alleles. Some archaic TCR alleles, however, were distinct from alleles commonly found in modern human populations. We identified likely introgression candidates and investigated if they were nested within extended archaic-derived haplotypes. The identification of putative introgressed alleles was performed stepwise to eliminate variants that were ancestral to both modern and archaic populations, and variants that may have arisen through convergent evolution of single nucleotides within a gene. Given that both Neanderthal and Denisovan introgression likely occurred among modern human populations outside of Africa, we first identified the candidate allelic variants that were present in European or Asian populations and absent, or at very low frequency, in African populations. Second, to determine the extent of the archaic derived introgressed haplotype containing the TCR allele in question, we identified individuals from the 1KGP dataset that contained the same alleles and utilized its extended genomic sequences to leverage against the reference Neanderthal and Denisovan genomes. This allowed us to identify the specific SNP variants and haplotype length of the putative archaic introgressed segments in the TCR region of each of the individuals.

The introgressed regions around these alleles were then validated using several approaches: quantifying the variation between groups using fixation index calculation (F_ST_), identification of haplotype SNP motifs, summarizing using principal component analysis (PCA), and application of an approach that uses long-range linkage disequilibrium (sPrime) (32). We found four TCR alleles that were shared between non-Africans and the Vindija Neanderthal: TRAV12-2*02_S6060, TRAJ24*02_S1049, TRAJ26*01_S5236, and TRGV4*02_S0072 (**Figure 5A, Figure 5B, SI Figure 6A**).

**Figure 5.**
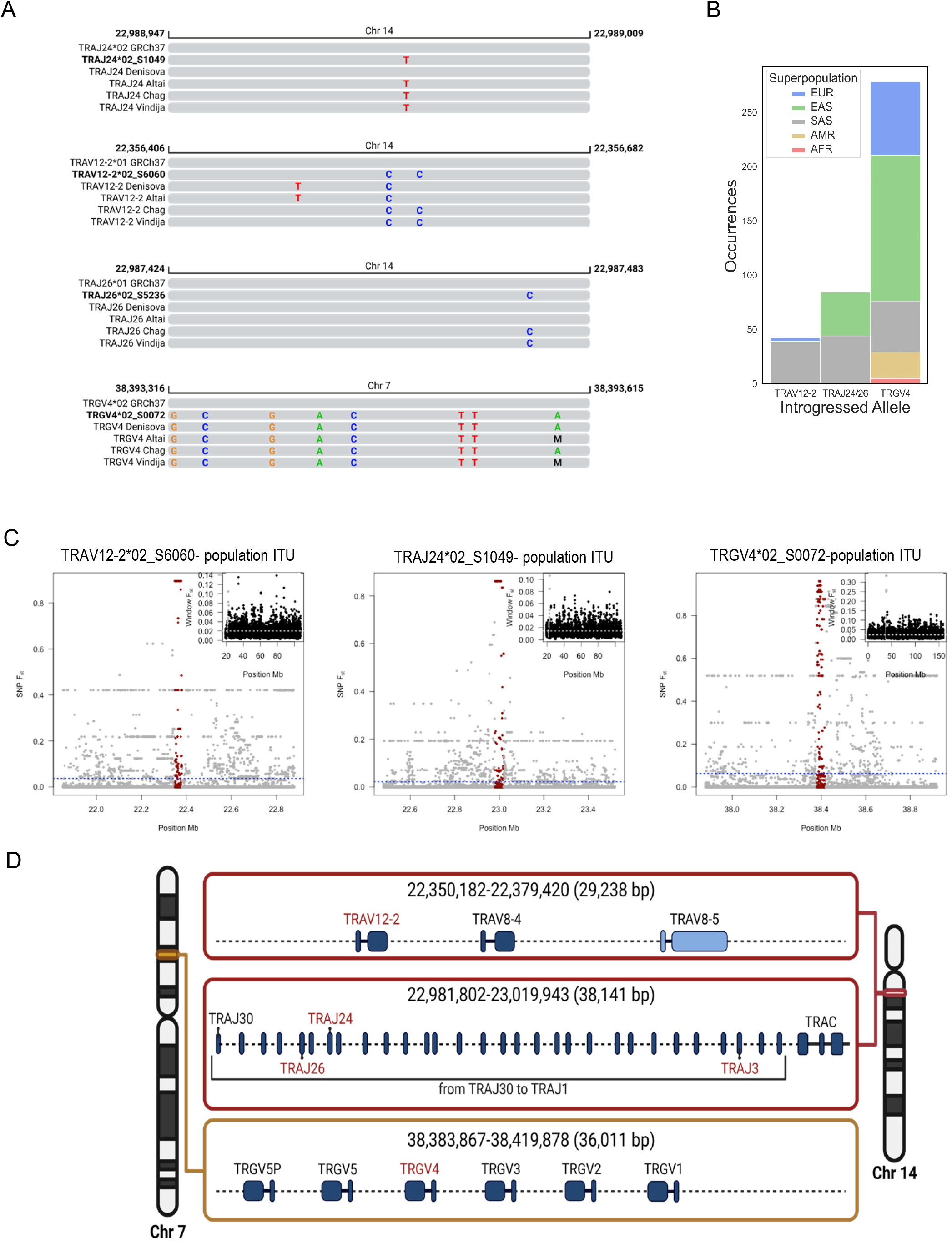
**A.** Alignment of novel TRAV and TRGV alleles with one Denosovan and three Neanderthal archaic and one modern reference assemblies. Upper panel left, alignment of novel allele TRAJ24*01_S1049 with consensus sequence spanning this gene from four archaic genome assemblies derived from one Denisovan individual and three Neanderthal individuals (Altai, Chagryskaya and Vindija) mapped to the GRCh37 reference assembly. The chromosomal coordinates are shown on the top bar for each panel. Nucleotide differences from the GRCh37 sequence are indicated in each alignment. Upper panel right, alignment of novel allele TRAJ26*01_S5236. Lower panel left, alignment of novel allele TRAV12-1*01_S6060. Lower panel right, alignment of novel allele TRGV4*02_S0072. **B.** Occurrences of Introgressed Alleles in 1000 Genomes Super Populations. Stacked barchart illustrating the frequency of each of the introgressed alleles in the five 1000 Genomes population groups. The population groups key is shown within the panel. **C**. Chromosome-wide Windowed F_ST_ values for the 1000 Genomes Indian Telugu in the UK (ITU) population as an illustration. The ITU population was split based on whether individuals had the candidate introgressed allele in question or not, and fixation indices were computed between these two groups. Values were averaged over 50kbp windows with 25kbp steps and the dots corresponding to the introgressed region were colored in red. **D**. Introgressed haplotypes identified in the study. Illustration of the two TRA genomic regions identified as archaic origin introgressed segments. The upper segment comprises a 29.2 kb segment that includes the functional TRAV12-2 and TRAV8-4 genes and the TRAV8-5 pseudogene, the middle segment includes a 38.1 kb region encompassing the TRAJ30 to TRAC genes, both segments located on chromosome 14. The lower segment encompasses a 36 kb region of chromosome 7 including all TRGV genes between TRGV5P and TRGV1.

F_ST_ values were calculated between subgroups of 1KGP populations, separated by presence or absence of the putative archaic introgressed alleles. The introgressed regions, as identified by F_ST_ peaks of over 0.8 (**Figure 5C-D**), in the 1KGP GRCh37 VCF were the following: chr14:22981802 - 23019943 (~38kbp) for TRAJ24/26, chr14:22350182 – 22379420 (~29kbp) for TRAV12-2 and chr7:38383867 - 38419878 (~36kbp) for TRGV4. The TRAV12-2*02_S6060 allele was found in 42 1KGP individuals, TRAJ24*02_S1049/TRAJ26*01_S5236 in 84 1KGP individuals, and TRGV4*02_S0072 in 278 1KGP individuals. All TRAV12-2 and TRAJ24/TRAJ26 individuals were non-Africans and all but 5 (273/278, 98.2%) of the TRGV4 individuals were non-African (**Figure 5B**).

Using the Yoruban 1KGP population as the outgroup, Sprime identified an introgressed region overlapping the putative introgressed alleles in two non-African populations for the TRAV12-2 allele with a median of 195 SNPs over ~432kbp, two non-African populations for the TRAJ24/26 alleles with a median of 46 SNPs over ~126kbp, and 20 non-African populations for the TRGV4 allele with a median of 322 SNPs over ~1Mbp. It is likely that the TRGV4 signal is easier to detect due to the underlying haplotype being more common in several populations. We next determined the percent SNP calls in Sprime segments that had matching calls in the Neanderthal and Denisovan genomes. Sprime was executed on each 1KGP population separately, so the percentages were averaged across results from multiple populations. For the introgressed region containing TRAJ24/26, the percentage matching SNP calls was 70% and 17% for the corresponding Neanderthal and Denisovan regions, respectively. Two of the four populations where Sprime identified an introgressed region overlapping TRAJ24/26 were African, but the segment for both these populations had less than 15% matching SNP calls to either of the archaic genomes, suggesting that they may be false positives. The TRAV12-2 and TRGV4 segments had Neanderthal match percentages ranging from 50% to 60% and Denisova match percentages between 15% and 30%. Two of the TGV4 segments were found in African populations, but they had less than 5% matching SNP calls to the Neanderthal and Denisovan genomes. If Sprime positions that were not present in the Neanderthal were excluded, the Neanderthal match percentage increased to 83% for TRAJ24, 93% for TRAV12-2 and 86% for the TRGV4 region, providing evidence for a Neanderthal origin tract.

We identified SNP motifs for each of the putative introgressed segments by taking the SNPs whose most common nucleotide had a frequency difference of over 0.4 between the individuals with and without the alleles of interest within a 1KGP population and filtering out SNPs that were not in common with the Vindija. The TRAV12-2 region has a 22 SNP motif, the TRAJ24/TRAJ26 region has a 35 SNP motif, and the TRGV4 region has a 15 SNP motif. The TRAV12-2 introgressed allele is most frequent in modern South Asian population, TRAJ24 in both South and East Asian populations and the TRGV4 introgressed allele is present in all non-African populations with the highest frequency found in East Asian and Europeans (**Figure 5B**). The TRAJ24*02_S1049 and TRAJ26*01_S5236 alleles immediately appeared to be part of the same putative introgressed region since they were found in the same study donors and are both present in the same set of 1KGP individuals. The alleles are identifiable as part of the same haplotype in donor D14 (**SI Figure 6B**). The most frequent introgressed TCR region, the TRGV4*02_S0072 containing haplotype, was clearly identifiable within a subset of the Southern Han Chinese population using principal component analysis (PCA) (**Figure 6A**). The introgressed TRGV4*02_S0072 allele contains eight nucleotide differences compared to the closest known TRGV4 allele, TRGV4*02, resulting in four amino acid changes that alter the CDR1, CDR3 and HV4 regions of the V gene (**Figure 6C-D**).

**Figure 6.**
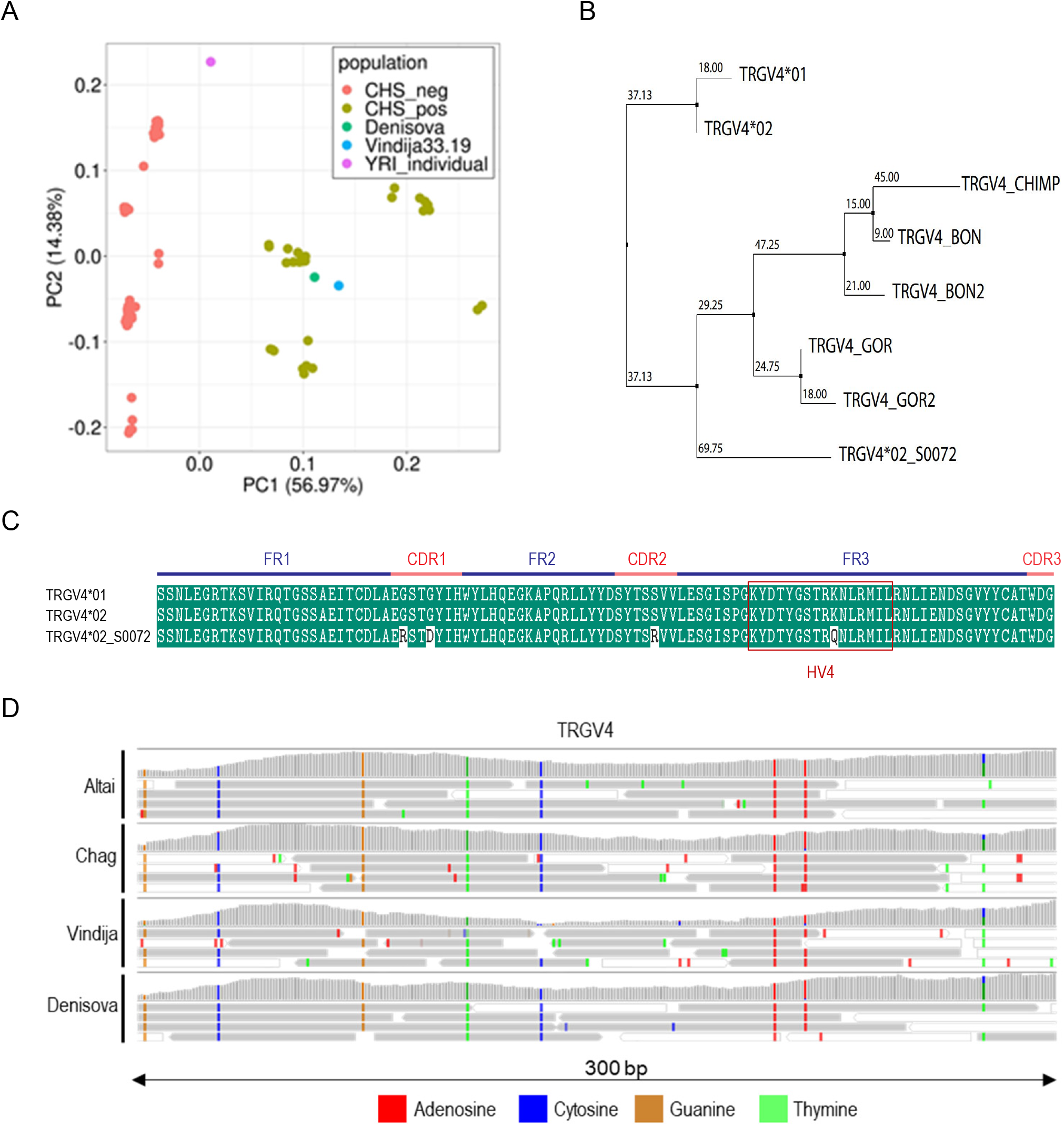
**A.** Principal Component Analysis of the TRGV4*02_S0072 introgressed segment for CHS. The TRGV4*02_S0072 introgressed region was defined as chr7:38383867-38419878 according to the region found by Sprime. CHS_pos individuals are defined as the members of the Southern Han Chinese (CHS) population set who contain the TRGV4*02_S0072 allele, while CHS_neg individuals are those who do not have it. **B.** Phylogenetic tree of TRGV4 alleles. Human alleles, TRGV4*01, TRGV4*02 and introgressed allele TRGV4*02_S0072 are shown in addition to TRGV4 alleles in Bonobo, Chimpanzee and Gorilla. **C**. Alignment of the translated protein sequence of previously known TRGV4 alleles with the candidate introgressed TRGV4*02_S0072 allele. The upper bar shows the positions of functional framework, CDR1, CDR2 and HV4 regions of the TRGV4 gene to illustrate the positions of amino acid variation specific to the introgressed allele. **D.** Illustration of short read mapping of archaic DNA reads from TRGV4 candidate introgressed genes. BAM files from the single Denisova assembly and three Neanderthal assemblies, Altai, Chagryskaya and Vindija, were used to visualize the short-read mapping using the IGV (The Integrative Genomics Viewer) genomic viewer. The nucleotide difference to the GRCh37 assembly is shown, the color key to the archaic specific variant nucleotides is in the key below the figure. The bar chart above each short-read mapping picture shows the consensus of all the short reads at each position of the gene.

Finally, comparative analysis of the TCR genes present on the available reference assemblies of Bonobo, Chimpanzee, Gorilla and Rhesus macaque was performed. A total of 193, 190, 221 and 265 TCR alleles were identified for each species respectively, presumably reflecting the overall degree of genomic coverage of the TCR loci. The analysis of multiple assemblies for each species reveals a degree of allelic diversity amongst multiple genes, and the presence of non-functional allelic variants, similar to that identified amongst human subjects. As shown in figure 7 and supplemental figure 7, The variations were primarily located in the CDRs, with the framework regions being highly conserved between all primate species examined (**Figure 7A-B, SI Figure 7**). Human TRAV14 alleles, for example show a P/Q amino acid variation in the CDR1 region, similar to that found in two chimpanzee TRAV14 alleles, whereas variation in the CDR3 region of TRAV14 is found in chimpanzee, bonobo and rhesus macaque TRAV14 (**Figure 7A**).

**Figure 7.**
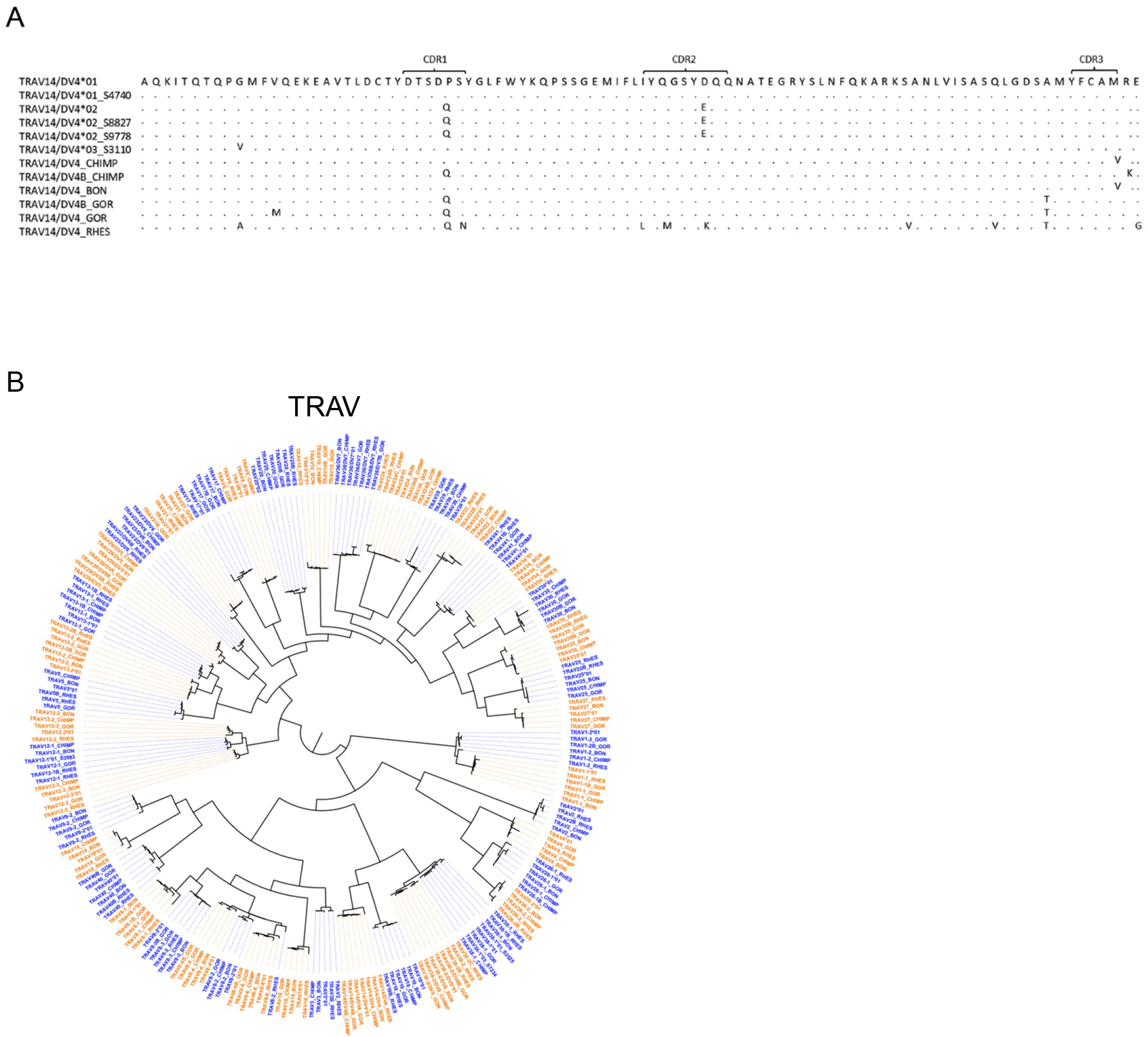
**A**. Alignment of the amino acid translations of all TRAV14 alleles from human, Bonobo, Chimpanzee, Gorilla and Rhesus macaque. **B**. Circular dendrogam of TRAV alleles of Bonobo, Chimpanzee, Gorilla and Rhesus macaque alongside the closest human allele for each TRAV gene.

## Discussion

Understanding variation within components of the human immune system is important for studies of disease susceptibility (33), prognosis (34) and population history (35). Most of the genetic variation identified to date has been ascribed to the MHC loci (36). However, polymorphisms in the genes that encode the TCR, the molecules that bind antigenic peptides in the context of MHC molecules, has not been viewed as a critical component of this process. Only low levels of variation within the TCR locus have been reported to date and the reference database of TCR gene variation, the IMGT TCR database contains low numbers of alleles for almost all TCR genes although recent evidence involving a study of TCRB genes reveals additional variation may be plentiful (19). The current study reveals, however, far higher levels of diversity in expressed TCR VDJ genes than previously reported. We describe almost double the numbers of TCRV alleles previously identified and a 30% increase in the total number of TRAJ alleles.

In contrast to the immunoglobulin heavy chain (IGH) locus (37), we found that genomic deletions are relatively infrequent in the TCR loci. One previously described common deletion region encompassing TRBV4-3 was identifiable here through loss of expression of this gene in multiple individuals. Homozygous genomic deletion encompassing TRGV4 and TRGV5 was shown for one case (**SI Figure 3B**). Inferred haplotype analysis was used to validate novel alleles identified in this study and to identify frequent hemizygous loss of expression. Our data revealed that many of these were not due to genomic loss but due to the presence of allelic polymorphisms that resulted in low expression of non-functional alleles, ENFs. The ENFs were characterized by SNPs that result in stop codons, likely leading to unstable mRNA transcripts. Three TRAV ENF alleles were identified in the donor cases, TRAV1-2*03_S6094, TRAV19*01_S1366 and TRAV34*01_S2006, and four TRBV ENF alleles, TRBV10-1*02_S3492, TRBV30*02_S9206, TRBV6-8*01_S7864 and TRBV6-8*01_S2439. TRBV10-1*02_S3492 was found in 14/45 (31%) individuals from multiple populations and therefore can be expected to be homozygous in a significant percentage of the human population.

The presence of non-functional allelic variants of TCR genes at such levels is suggestive of a mosaic functionality in the TRA and TRB loci. If this is a common feature of TCR genotypes, we could expect to see non-functional allelic variants in closely related species. Indeed, a comparison of the TCR genes present in the available haploid genomic assemblies of the closest living species to humans, Chimpanzee and Bonobos, reveals a total of six TCR V genes: TRAV38-1, TRBV10-3, TRBV11-2, TRBV4-1, TRBV5-4 and TRBV9, with allelic variants containing stop codons, in addition to variants without stop codons present in other assemblies of the same species, such as TRBV14 in Gorilla or TRBV7-2 in Rhesus macaque.

An additional source of allelic diversity that affects TCR gene function was identified with splice site and RSS site SNP variations that correlated with loss of expression in particular populations. Splice site variations were identified in TRAV26-2 and TRAJ12 and RSS variation in TRAJ17. For each of these alleles the frequency of the SNP variant within the different 1KGP groups corresponded closely with the frequency of hemizygous loss of expression found in the donor cases analyzed here from the same populations. Loss of TCR gene expression due to common polymorphisms that result in loss of function varied in frequency between different populations. However, differences in the frequency of functional alleles remained the most common factor underlying inter-individual and inter-population variation. TCR allelic diversity was highest in sub-Saharan African donors for most TCR genes, a finding that is consistent with multiple studies of genetic diversity amongst human populations (25), with 30, 33, eight and five unique alleles in TRAV, TRBV, TRGV and TRAJ genes, respectively. TRDV exclusive alleles showed little variation in frequency between different population groups.

The TCR response to the immunodominant influenza M1 epitope was shown to be restricted and include a public TCR that uses TRBV19 and TRAV38/J52 (38). The presence of a common ENF allele of TRAJ52 in East Asian populations would inevitably prevent the formation of such TCRs in homozygous individuals. In the same population TRAJ17 expression is frequently lost in hemizygous or homozygous form due to an AAGT deletion variant (rs3841038), which removes the TRAJ17 splice donor site. This TRAJ gene was found to be the most common TRAJ gene used in Gag293-specific TCRs isolated from HIV-1 controller cases (39). The TRAJ12 gene, along with TRAJ20 and TRAJ33, is frequently found utilized in a restricted TRAV1-2 using TCR in mucosal-associated invariant T (MAIT) cells (40, 41). Our results show that the frequent loss of expression of TRAJ12 in African individuals is due to SNP variant A/G rs62622786, which removes the RSS of TRAJ12. Individuals homozygous for this non-functional TRAJ12 allele are common among African individuals, which has implication for MAIT cell responses in this population group. As shown by these studies and several other reports (42), germline-encoded variations affecting CDRs may be especially important for shaping the T cell repertoire.

In the current study, we found frequent variations mapping to CDRs. Five TRAV genes had novel coding variants in the CDR2, and three variants displayed coding changes at the CDR2-FR3 border. We further identified variants of TRAV12-1 and TRAV8-2 with coding differences within the CDR3 segment at the 3’ end of the V gene, and CDR1-located coding variation in two novel alleles of TRAV38-1. We observed even more frequent coding changes due to TRAJ allelic differences, with six novel TRAJ alleles having coding variants within the CDR3 segment of those genes. Three TRBV genes with coding variation within the CDR1 was also identified, and one allele, TRBV18*01_S1676, with a coding difference at the FR2-CDR1 border. Four TRBV genes had novel alleles with coding changes within the CDR2 and one with a variation at the CDR2-FR3 border. Two novel TRBJ variants, TRBJ1-4*01_S8217 and TRBJ2-3*01_S5123, show coding changes within the CDR3. For TRGV we identified four novel alleles containing coding variation within these regions. TRGV3*01_S0606 contained a CDR2 coding variant and TRGV4*02_S3970 contained a coding change within the CDR1. Two novel TRGV4 alleles, TRGV4*01_S0072 and TRGV4*02_S2864 contained coding variation within both the CDR1, CDR2 and the hypervariable 4 (HV4) segment of FR3.

The presence of coding variation within the CDR3 of several novel alleles enabled the identification of these alleles in VDJdb, a curated database of published TCR CDR3s (43). For example, TRAV47*01_S5950, was identified in multiple published TCR CDR3 sequences. It is involved in HLA-A*02-restricted presentation of the Influenza A M epitope GILGFVFTL, HLA-A*02-restricted presentation of the CMV gene p65 epitope NLVPMVATV, and HLA-B*44-restricted presentation of CMV gene IE2 epitope NEGVKAAW. TRBJ1-4*01_S8217 is involved in HLA-A*02 associated binding to CMV p65 epitope NLVPMVATV and HIV Gag-1 epitope RTLNAWVKV and in HLA-A*03 associated binding to CMV IE1 epitope KLGGALQAK. Furthermore, TRAJ2-3*01_S5123 can be identified in a TCR that recognizes an HLA-A*02-restricted influenza A M epitope GILGFVFTL.

TRB V and J usage was shown to be useful in guiding prognosis and immunotherapies in head and neck cancer cases, with dominant expression of specific V and J genes correlating with survival rates both in individuals with specific HLA allelic backgrounds and independent of HLA status (44). Intratumoral γδ T cells are the most favorable prognostic indicator in multiple cancers (45). TCR variation is also important in autoimmunity. A study of the potential role of the influenza Pandemrix vaccine in inducing POMT1 autoantigenic responses identified public TRAV10/TRAJ24 CDR3s in narcolepsy cases that are specific for TRAJ24*03 (46) rather than any of the other three TRAJ24 alleles. As allelic variation of MHC alleles is known to affect predisposition, protection from and prognosis of multiple cancers and autoimmune disorders, TCR germline gene variation impacting this interaction will likewise be important.

TRGV allelic variation is large. We identified a total of eight allelic variants of TRGV4, several of which were present in contrasting frequencies in different populations. One novel allele identified in this study, TRGV4*02_S0072, has eight nucleotide differences from TRGV4*02, found in the GRch37/hg19 assembly. It is found almost exclusively in non-African Eurasian populations. We found an identical sequence in three Neanderthal high coverage assemblies and additionally in the Denisovan assembly. We validated this allele using targeted genomic PCR and Sanger sequencing. Furthermore, analysis of 1KGP samples containing the same allele allowed us to delineate an extended haplotype encompassing this allele in these cases, revealing an introgressed region of 36 kb present in multiple individuals. The presence of this allele at high frequency in Eurasian populations suggests the possibility of this being an adaptive variant. The recent evolutionary history of human populations can also underly allelic frequency variations affecting disease susceptibilities. Two loci that contain variants introduced to the modern human gene pool through Neanderthal introgression events were shown to modulate the response to Covid-19. One locus was present on chromosome 3, containing the CXCR6 and CCR9 chemokine receptor genes, which increased the risk of severe disease(47). The other was on chromosome 12, containing the OAS1–OAS3 genes, which reduced the risk of developing severe disease(48). The TRGV4*02_S0072 allele contains four coding variations, two within CDR1, one within CDR2 and one within the HV4 loop region. TRGV4 germline sequences were recently shown to be critical, and mutations introduced within the HV4 loop strongly inhibited BTNL3 binding (49). BTNL3 is expressed in gut epithelial cells and in combination with BTNL8 is an important regulator of Vg4+ T cells. This subset of γδ T cells are members of the intestinal intraepithelial lymphocyte (IEL) cell population, which are involved in processes that modulate inflammation and protection from infection (50). The importance of the BTNL3/BTNL8 Vg4+ interaction was highlighted by Mayassi et al. 2019, who found that gluten induced inflammation in Celiac disease resulted in a disruption of BTNL8 expression and subsequent disruption of Vg4+ T homeostasis (51). The presence of multiple TRGV4*02_S0072 coding variations in the CDR1, CDR2 and HV4 loops, critical for BTNL3 binding suggest that the presence of this allele may play an important role in this regulatory pathway. If loss of function of TRGV4 is advantageous in some environmental circumstances, then alternative means of disrupting the TRGV4/BTNL3 pathway may be relevant. We identified homozygous deletion of TRGV4 in one case (D12) in our study (**SI Figure 3B-C**). The BTNL8/BTNL3 region itself is the location of a 56 kb genomic deletion (52) that disrupts both genes and is found at highest frequency in European, East Asian, and American populations and lowest in African, Middle East and Oceanic populations. The conclusion that TRGV4*02_S0072 is present in an introgressed haplotype was strengthened by comparison of this allele with the TRGV4*01 and TRGV4*02 alleles and allelic variants of the gene found in Chimpanzee, Bonobo and Gorilla. These great ape TRGV4 allelic variants were found to show a closer identity to TRGV4*02_S0072 than to TRGV4*01/2, indicating that the former allele may have evolved within archaic human ancestors for a substantial period prior to the introgression event (**Figure 6B**). In contrast, the Neanderthal derived K66->Q66 difference in TRAV12-2*02_S6060 appears to be more recent. Both Gorilla and Rhesus macaque have a lysine at that codon, resembling the five remaining TRAV12-2 alleles we found. Both Bonobo and Chimpanzee, however, have a glutamic acid codon at this position, indicating convergent change to the same residue in both neanderthal and Chimpanzee/Bonobos.

The results of this study indicate that genetic diversity in the TCR locus is extensive, both in terms of individual variation and at a population level. The origin of this diversity and population-based diversity various demographic forces such as mutation, natural selection, population bottlenecks and gene flow between modern human populations between modern and archaic human populations (53, 54). We show that the number and diversity of V alleles is vastly under-represented in the current public reference database and demonstrate that most of the variation in the newly discovered V and J alleles results in coding changes many of which involve the CDR1, CDR2, CDR3 and HV4 regions of TCR genes and may affect functionality of the genes. Allelic diversity in the TCR loci was apparent in all other primate assemblies that we compared to the compiled human database. The human TCR is important in defense against pathogens, autoimmunity, tissue regulation and cancer and the results provided in this study will help define critical components that may have been hitherto overlooked.

## Figure legends

**SI Figure 1**

Allelic variation in TRAJ and TRBJ genes and position of coding changes in novel V and J alleles. **A.** Circular dendrogram illustrating the TRAJ alleles identified in the 45 cases analyzed. Genes are alternated in blue and red color to allow easier visualization. Novel alleles are indicated by the presence of a suffix consisting of an _S and 4 numbers that are products of the IgDiscover inference program. **B.** Circular dendrogram illustrating the TRBJ alleles identified in the 45 cases analyzed. **C.** Circular dendrogram illustrating the TRDV alleles identified in the 45 cases analyzed. **D**. Upper pie chart, illustration of the numbers of TCR V gene allelic variants in the 45 donors that result in a codon difference compared to known alleles. Lower pie chart, illustration of the proportion of coding variants identified in novel TCR J genes expressed in the 45 cases. **E.** Alignment of the translated amino acid sequence of the TRAJ alleles. Novel TRAJ allele amino acid changes are highlighted in green. **F.** Alignment of the translated amino acid sequence of the set of novel TRBJ alleles. Novel TRBJ allele amino acid changes are highlighted in green. The J gene CDR3 and framework 4 regions are indicated on the top of panels E and F.

**SI Figure 2**

Validation of novel alleles identified. **A.** Numbers of alleles validated by different measures. These evidences are: present in multiple individuals; present in both individuals of a set of monozygotic twins; present in non-twin familial cases; present in both twin and familial cases; genomically validated following PCR and Sanger sequencing using from DNA from a 1000 genomes sample case; genomically validated following PCR and Sanger sequencing from one of the 45 donors in the current study; genomically validated following PCR and Sanger sequencing in both a donor case and, separately a 1000 genomes sample case DNA sample; present heterozygously in the output of IgDiscover *plotalleles* module haplotype analysis; present hemizygously in the output of IgDiscover *plotallele*s module haplotype analysis, and present homozygously in the output of IgDiscover *plotalleles* module haplotype analysis. The barchart on the left shows the numbers of novel alleles validated within the various categories. **B.** IgDiscover haplotype analysis of the TRAV locus in a three-member family group, comprising two parents, D45 and D25 and an offspring, D05. In all three cases the TRAJ gene TRAJ13 was heterozygous for the *01 and *02 alleles and therefore suitable as a haplotyping anchor. The dotted blue lines on the right of the haplotype panels indicate the parental origin of the inherited TRAV chromosomal haplotypes of D05.

**SI Figure 3**

TRDV, TRGV and TRAJ gene content variation between individuals **A.** Illustration of TRDV and TRADJ variation at the gene level. Inferred haplotype analysis identified the presence of homozygosity, heterozygosity, or deletion that are denoted by blocks of color, the key being present on the top left above the figure. The population groups of the case are shown below the figure, the key being present on the top right of the figure. For each case the gene names are on the right-hand side in chromosomal order and the case numbers are shown below the figure. **B.** Illustration of variation in the TRGV locus at the gene level. Cases that were informative for haplotype analysis or by lack of TRGV expression are shown. **C.** Genomic analysis of TRGV status. Illustration of the result of a multiplex PCR was performed on a set of donor DNA samples using a primer set mix that simultaneously amplified parts of four TRGV genes, namely TRGV9, TRGV4, TRGV2 and a control target in a separate genomic region, namely HOX3. Sample M=DNA size marker, NTC=non-template control. The bands corresponding to each of the four targets are indicated on the right of the gel picture. **D.** Illustration of TRAJ variation at the gene level. Inferred haplotype analysis identified the presence of homozygosity, heterozygosity, hemizygous, or homozygous presence that are denoted by blocks of color, the key being present on the top left above the figure. The population groups of the case are shown below the figure, the key being present on the top right of the figure. For each case the gene names are on the right-hand side in chromosomal order and the case numbers are shown below the figure.

**SI Figure 4**

Haplotype analysis of the TRAJ locus reveals selective loss of TRAJ expression. The upper analysis shows the result of IgDiscover *plotalleles* haplotype analysis of the TRAJ locus in case D34 haplotyped using the heterozygous anchor gene TRAV8-2 showing TRAJ12 loss from one chromosome. The lower analysis shows case D28 haplotyped using the heterozygous anchor gene TRAV8-3 showing TRAJ17 loss from one chromosome. **B.** Schematic of genomic variation at the borders of the TRAJ genes TRAJ12 and TRAJ17. The positions of the RSS heptamer and splice donor site of both genes are shown in addition to the position of the SNP variants rs62622786 (G/A) and rs3841038 (AAGT/−) marked in red. In each case the frequency of each variant in the 1000 genomes population sets are shown in the lower pie charts. **C.** Association of heterozygous TRAJ52*01_S5131 alleles with different chromosome to TRAJ52*01. Left panel, Case D02 is heterozygous for TRAV26-1*01 and TRAV26-1*02. These TRAV alleles recombine locally with TRAJ52 alleles on the same chromosomal locus showing that TRAJ52*01 is present on the same chromosome as TRAV26-1*02 while TRAJ52*01_S5131 is present on the TRAV26-1*01 containing chromosome. Right panel, Case D27 is heterozygous for TRAV4/DV4. TRAJ52*01 is present on the same chromosome as TRAV14/DV4*01_S4740 while TRAJ52*01_S5131 is present on the TRAV14/DV4*02 containing chromosome.

**SI Figure 5**

**A.** Haplotype analysis of TRAV in case D31 using the IgDiscover *plotallele* module. Counts of TRAV alleles recombined with haplotying anchors TRAJ44*01 and TRAJ44*01_S9876 and shown in chromosomal order. Novel alleles present homozygously are denoted with a red asterisk and those present heterozygously denoted with a blue asterisk. **B**. Genomic validation of novel alleles using BAC sequence analysis. Illustration of BAC clones showing 100% sequence identity to novel and known TCR alleles are shown with the sequential positions of functional TCR genes marked with a vertical orange bar. The allele name is shown above each gene position and the novel alleles are marked with a horizontal red bar. The top five BAC clones; BACs RP11-114B6, RP11-122A11, RP11-654L23, RP155E15 and RP11-325O4, contain sequence from the human TRBV locus and the lower BAC, CH17-155D14, encompasses the TRGV region.

**SI Figure 6**

**A.** Illustration of short read mapping of archaic DNA reads in each of the candidate introgressed genes. BAM files from the single Denisova assembly and three Neanderthal assemblies, Altai, Chagryskaya and Vindija, were used to visualize the short-read mapping using the IGV (The Integrative Genomics Viewer) genomic viewer. From top to bottom, the panels show the TRAJ24 gene, the TRAJ26 gene and the TRAV12-2 gene. In each case the nucleotide difference to the GRCh37 assembly is shown, the color key to the archaic specific variant nucleotides is in the key below the figure. The bar chart above each short-read mapping picture shows the consensus of all the short reads at each position of the gene. **B**. Haplotype analysis of case D14 using the IgDiscover *plotalleles* module, using heterozygous anchors from the gene TRAV8-3 to separate the expressed TRAJ alleles in this case. The TRAJ alleles are shown in chromosomal order. The grey bar on the bottom shows the part of the TRAJ region present within the candidate introgression segment in D14.

**SI Figure 7**

Circular dendrogams of TCR alleles of Bonobo, Chimpanzee, Gorilla and Rhesus macaque alongside the closest human allele for each TCR gene. **A.** TRAJ, **B.** TRBV, **C.** TRBJ, **D.** TRGV, **E.** TRDV, **F.** TRGJ, **G.** TRDJ, **H.** TRBD and **I.** TRDD.

## Acknowledgements

We thank the blood donors and Dr. Anton Sendel for collection of samples. We also thank Jonathan Coquet and Xaquin Dopico Castro for helpful input on the manuscript, and Fondation Dormeur, Vaduz for generous support of equipment.

## Author Contributions

Conceptualization: MCo and GBKH

Methodology: MCo, MCh, MK and MJ

Investigation: MCo, MM, MCh, SN, MK, CB, ML

Visualization: MCo, MCh, SN and GBKH

Resources: CS, AF

Funding acquisition: GBKH

Supervision: MCo, MJ, GBKH

Writing – original draft: MCo, GBKH

Editing – original draft: MCo, MM, MCh, SN, MK, CB, ML, MJ, GBKH

## Human subjects and ethical statement

Peripheral blood mononuclear cells were isolated from whole blood samples collected with informed consent from healthy volunteer donors that enabled the donors to self-identify their population group (ethics permits #2016/1053-31 and #2017/852-32), and from a cohort of malaria-infected individuals from central Africa (ethics permits #2006/893-31/4, #2013/550-32) obtained from Regionala Etikprövningsnämnden, Stockholm. The samples were pseudo-anonymized in the study. The five sets of TRA and TRB twin libraries utilized in this study are publicly accessible sequence data from Rubelt et al 2016 (20).

## Method details

### RNA and DNA isolation

Peripheral blood mononuclear cells were isolated from whole blood samples using Ficoll-paqueTM (GE Healthcare Life Sciences), with subsequent RNA and DNA isolation as detailed in Vazquez Bernat et al. 2019 (55).

### TCR library preparation

TCR cDNA libraries were created using chain specific cDNA synthesis primers. Individual primers specific for the constant regions of TRA, TRB, TRG and TRD were utilized with 400 ng of total RNA per reaction, using the Sensiscript cDNA synthesis kit (Qiagen). This cDNA synthesis primer also includes a 21 nucleotide UMI that is present in the amplified product and is used in downstream computational analysis. The cDNA was purified using the Qiagen PCR purification kit and resuspended in 10 µl of H2O. TCR library amplification was performed separately for TRA, TRB, TRG and TRD using chain specific 5’ multiplex primer sets, along with a universal reverse primer that recognizes the amplification target site present in the 5’ tail of the chain specific cDNA synthesis primer. The TCR library PCR products were subsequently gel purified and indexed for sequencing using the index primer sets and conditions described in Vazquez Bernat et al, 2019 (55).

### Genomic validation

A set of novel alleles were validated by Sanger sequencing following target PCR amplification using primers located upstream of exon 1 and downstream of exon 2 of each V gene or targeting the genomic sequence encompassing a novel J gene.

### NGS sequencing and computational analysis

Individually indexed libraries were sequenced using the MiSeq V3 2 × 300 cycle system (Illumina). The R1 and R2 reads were processed using the IgDiscover (TCR version). In brief, the R1 and R2 sequence reads were merged, the UMIs were extracted, and the sequences were assigned to a starting database, consisting of the IMGT TCR reference database for each TCR chain. The IgDiscover program assigns each TCR sequence to the closest database reference sequence and calculates specific nucleotides that differ from this assignment. The identification of a consensus sequence that differs from a starting database sequence results in a potential candidate novel germline sequence. The program then uses a series of filters designed to identify features of germline sequences and to remove non-germline sequences from the set of candidate sequences. These filters, primarily based on the identification of multiple CDR3s or Js associated with candidate Vs or multiple CDR3s and Vs associated with candidate Js, function to identify unique V and J sequences that are utilized in multiple independent rearrangements. In addition, the program filters sequences for clonal expansion to handle expanded TCR clones can interfere with germline inference processes by obscuring heterozygous alleles of the same gene.

### Genotype analysis

The IgDiscover analysis program (TCR version) incorporates a feature designed specifically for genotypic analysis of Adaptive Immune Receptor content. The *corecount* module inputs the set of filtered sequences produced by IgDiscover. For each gene, unique UMI containing sequences identical to alleles in the database are counted. In contrast to the germline inference process the *corecount* genotypic analysis module enables identification of low expressed genes. *corecount* genotypic analysis was performed on V and J sequences from all libraries and D sequences from TRB and TRD libraries.

### Genotypic comparison

The individualized V, (D) and J genotypes for each case were collected for individual and population level comparison. The genotypic outputs enabled the identification of V, J allelic content for TRA and TRG and V, D and J content for the TRB and TRD loci. The haplotype results enabled the identification of gene level variation in cases where well expressed heterozygous D or J alleles could be used as anchors. Genes were scored as homozygous if the same allele was identifiable on both chromosomes (maternally and paternally derived). Genes were scored as heterozygous if a different allele of the same gene was present on each chromosome. Genes were scored as hemizygous if a single allele for a gene was present on one chromosome and no allele for the same gene was detectable on the other chromosome. Genes were scored as duplicated if more than one allele for a single gene was detected on a single chromosome.

### Analysis of TCR Introgression

We started by identifying TCR alleles that are exclusive to non-African populations in our dataset and that are shared with either reference Chagyrskaya, Altai, Vindija Neanderthal or Denisovan individuals. Next, we identified individuals in the 1KGP phase 3 VCF dataset which had these alleles. These individuals were labeled as “positive” if they had the introgressed alleles. We next performed Fixation Index calculations using vcftools 0.1.17 (56) between individuals with and without the introgressed alleles in a single 1KGP population. To perform an independent validation of the introgressed regions found by high FST values when comparing with and without introgressed alleles, we utilized Sprime (57). The S∗-like approach, Sprime, is a robust bioinformatic tool intended to find archaic introgressed segments in target populations without the need for a reference archaic genome. It relies on identifying distinct mutational characteristics of archaic introgressed segments in target populations, by leveraging it against an African outgroup who do not carry the archaic introgressed segments. The method finds the candidate introgressed regions that are sets of tightly linked SNPs and that are highly divergent to the African outgroup, by maximizing a pairwise score using a dynamic programming approach. We used version 07Dec18.5e2 of the S∗implementation described by Browning et al (57).

### SNP motifs

To find SNP motifs the introgressed regions have in common with the archaic humans, we first identified SNPs whose major alleles had at least a 0.4 frequency difference between the individuals who did or did not have the introgressed alleles within individual 1KGP populations. Next, we took the intersection between these high-frequency difference SNPs and the SNPs found in the Vindija genome.

### Principal Component Analysis

We computed PCA to verify that the individuals who carry the introgressed alleles cluster with the archaic individuals and separate from individuals who do not carry the archaic TCR haplotypes. The introgressed regions defined by F_st_ peaks were extracted from phase 3 (release 20130502) 1KGP VCF and subsequently merged with the Vindija and Denisovan VCF to perform PCA using the smartpca function of EIGENSOFT version 7.2.1 (58).

### Haplotyping

IgDiscover contains an inferred haplotype module named *plotalleles*. The module is based on the principle that heterozygous J or D alleles can be used to map associated V alleles to the appropriate chromosomal region. VDJ recombination occurs locally on a single chromosome and so a heterozygous J or D allele will only recombine with V alleles present on that chromosomal region and not on the corresponding locus of the other heterozygous chromosome. Hence maternally derived V alleles will associate with the maternally derived J or D allele and paternally derived V alleles will likewise only associate with the paternally derived J or D alleles on that chromosome. The *plotalleles* module creates a phased map of V and J alleles present based on the heterozygous J or D content of that case. In addition, heterozygous V alleles enable the identification of J or D gene deletion within the library. Haplotype analysis is utilized for two purposes in this study. First it is a useful validation process. The clear separation of a heterozygous novel germline allele serves as a means of validating the germline inference process. Secondly, the process enables the identification of structural or expression-based variation in each genotype, enabling the identification of duplications, deletions or loss of expression that may be due to recombination associated genomic variation. IgDiscover based inferred haplotype analysis was performed as described previously (22) using the appropriate heterozygous TCR anchors.

Zygosity inference was performed using a modified version of RAbHIT, Gidoni et al (59). Briefly, the method takes advantage of an allele’s co-occurrences with heterozygous “anchor” genes to calculate the probability of the co-occurrences (i.e. alleles that occur together in a VDJ rearrangement) given that the allele is on one chromosome or on two chromosomes. The Bayes factor (K) is the ratio of these two conditional likelihoods, and we consider lK = log10(K), which is positive when evidence favors homozygosity and negative when evidence favors heterozygosity. Sequences were preprocessed by filtering for <=2 V errors, 0 D errors, a <1E-4 D evalue, and 0 J errors. Next, the sequences were collapsed by unique V assignment, D assignment, J assignment, and CDR3 nucleotide sequence. In order to accumulate evidence from multiple anchors, we sum lK values across all anchors for a given allele, which is probabilistically appropriate given an independence assumption between rearrangements with different anchors. We used this approach to infer zygosity for all possible Vs, Ds, and Js. An lK with an absolute value of 3 was used as a cutoff to make a zygosity call. We implemented a system to find reliable anchors. For each individual, we found all pairs of heterozygous alleles that validated each other by providing an lK below −3 and used those as the anchors. In cases where there were no heterozygous Js to validate heterozygous Vs or vice versa, we found anchors by choosing the genes that had been confidently haplotyped by each other in other individuals, had a total minor/major allele ratio that was higher than the average for validated anchor pairs (.68), and were not commonly duplicated genes. The noise term was set to .104, which was determined by finding where the erroneous assignment rate of validated anchors plateaued with respect to the lK factor cutoff for heterozygosity.

### IgDiscover and *corecount* post-processing

For alleles in an individual which were not found by either *corecount* or IgDiscover, we required that the counts which were present to be over 20 to reduce false positives caused by IgDiscover inference. This criterium was not applied genes with low median frequency across all individuals compared to the alternate genes. These low expressed genes were defined as the five lowest median frequency TRAV genes (TRAV23, TRAV23/DV6, TRAV18, TRAV4, and TRAV40), five lowest median frequency TRBV genes (TRBV6-8, TRBV7-4, TRBV6-9, TRBV6-7, and TRBV10-1), and the two lowest median frequency TRGV genes (TRGV1 and TRGV5P). To reduce false positive end variants from IgDiscover inference, novel alleles which varied from known alleles only within the last eight nucleotides were required to have both *corecount* and IgDiscover results, since *corecount* would ensure presence of exact matches to the end variants. In addition, *corecount* results for high expressed genes (those whose mean frequencies are in the top 75% of their locus and gene type, TRAJ for example) were only taken if they have a count over five and frequency over 0.01%. TRBD *corecount* filtering required an allelic ratio of over 25%.

#### Software

Post-processing of IgDiscover results, Bayesian haplotype analysis, SNP analysis relating to 1KGP and introgressed regions, as well as generation of Figure 5B was performed using custom scripts in Python (version 3.7.4), utilizing the following packages: pandas (version 1.1.4), scipy (version 1.4.1), ensembl_rest (version 0.3.3), numpy (version 1.18.0), Biopython (version 1.79), scikit-allel (version 1.3.3), pyfaidx (version 0.5.9.2), gtfparse (version 1.2.0), pyVCF (version 0.6.8), and seaborn (version 0.11.0). The dendrograms were made using Aliview (version 1.26) and fasttree (version 2.1.11). Plots were made using R packages ggplot2 (version 3.1.0), gplots (version 3.0.1.1), ComplexHeatmap (version 2.9.3), dplyr (version 0.8.0.1) and ggvenn (version0.1.9).

#### Primate TCR gene comparison

Reference genome assemblies from Bonobo: panPan1.fa.gz, panPan2.fa.gz and panPan3.fa.gz; Chimpanzee: panTro3.fa.gz, panTro4.fa.gz, panTro5.fa.gz and panTro6.fa.gz; Western lowland Gorilla: gorGor3.fa.gz, gorGor4.fa.gz, gorGor5.fa.gz and gorGor6.fa.gz; and Rhesus macaque: rheMa2.fa.gz, rheMac3.fa.gz, rheMac8.fa.gz and rheMac10.fa.gz, were downloaded from Genbank. Individual primate sequences showing highest similarity to human TCR alleles were identified using the program BLAST.

