## Supplementary figures and images for "Archaic humans have contributed to large-scale variation in modern human T cell receptor genes"

### Supplemental figures

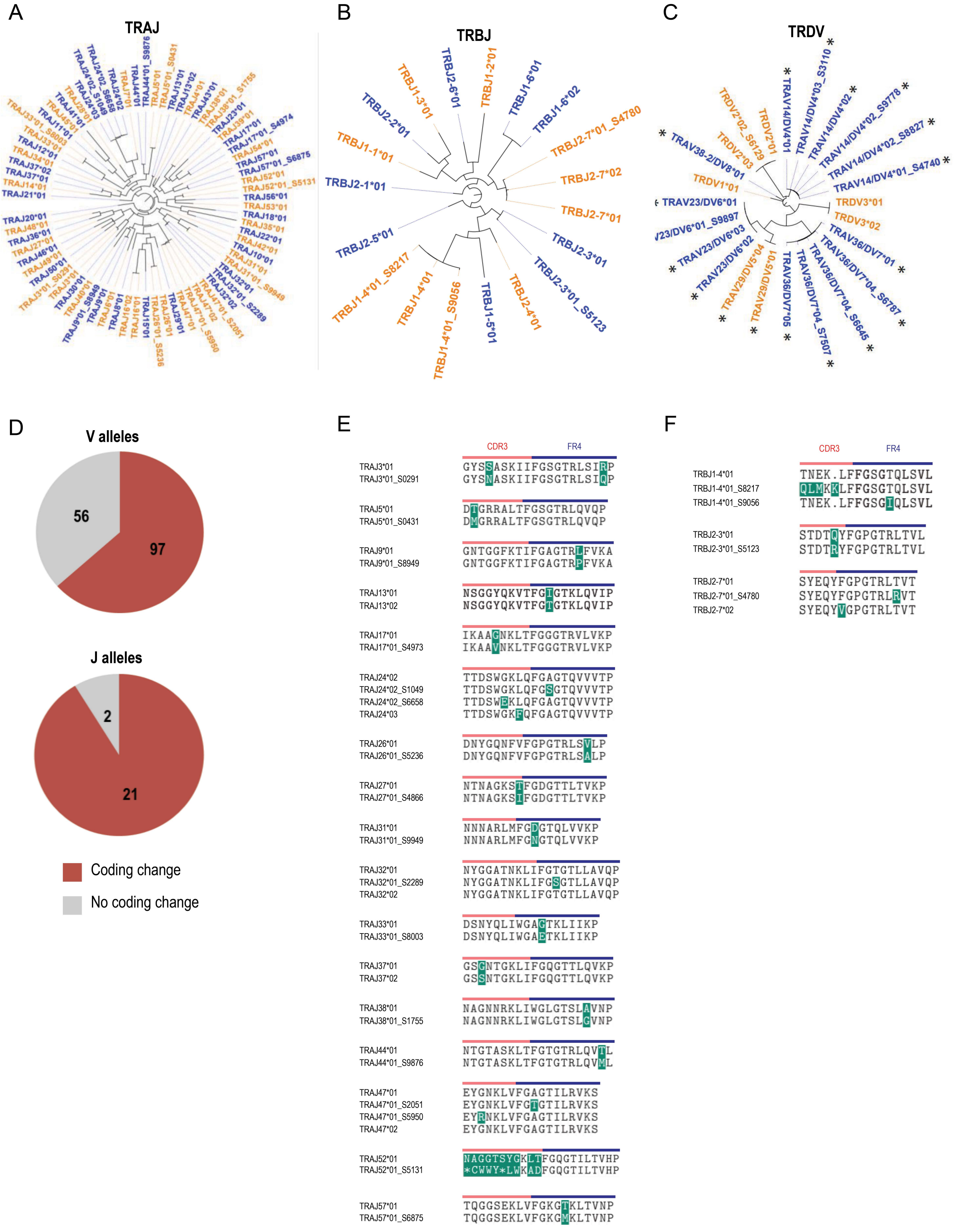

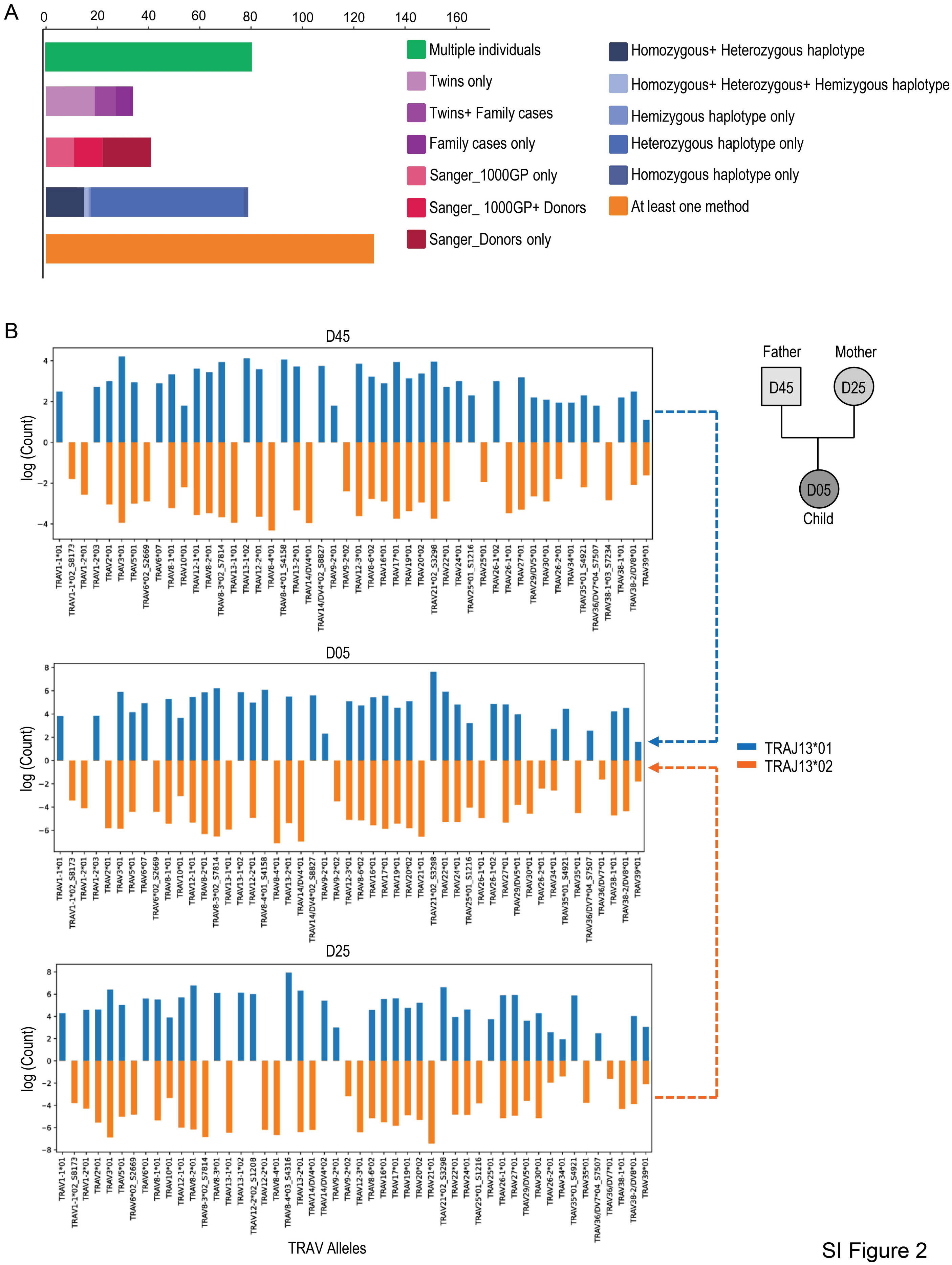

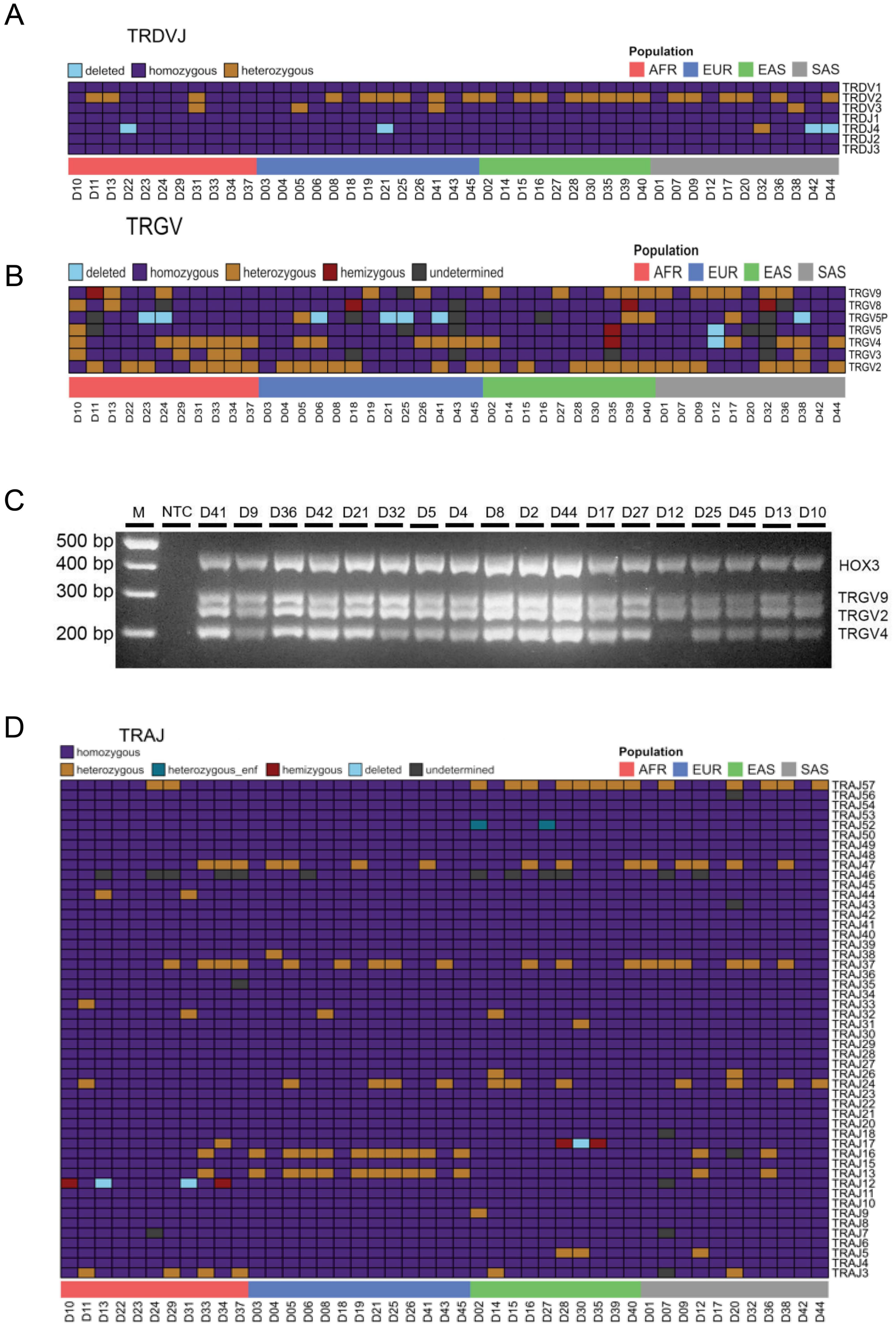

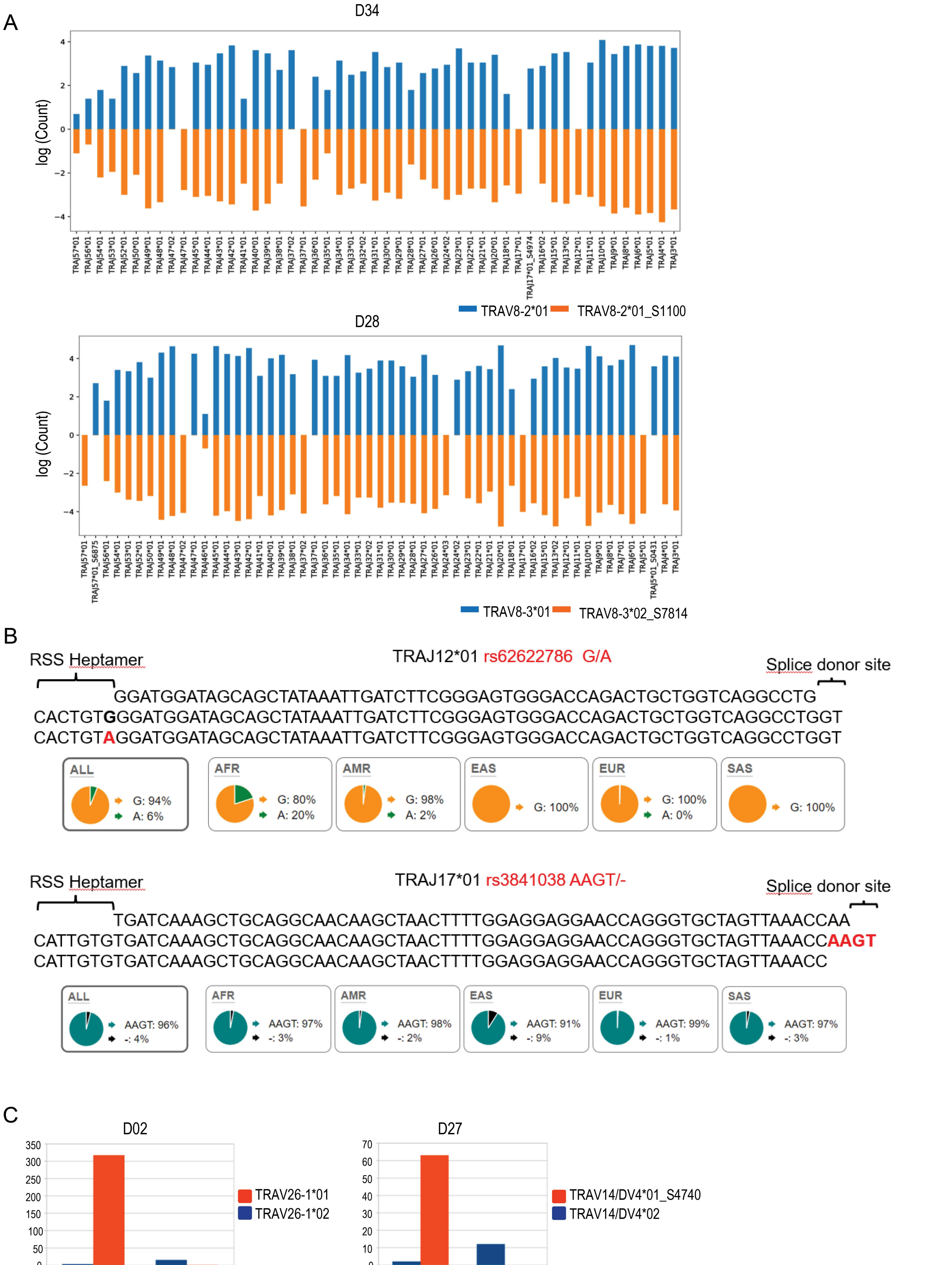

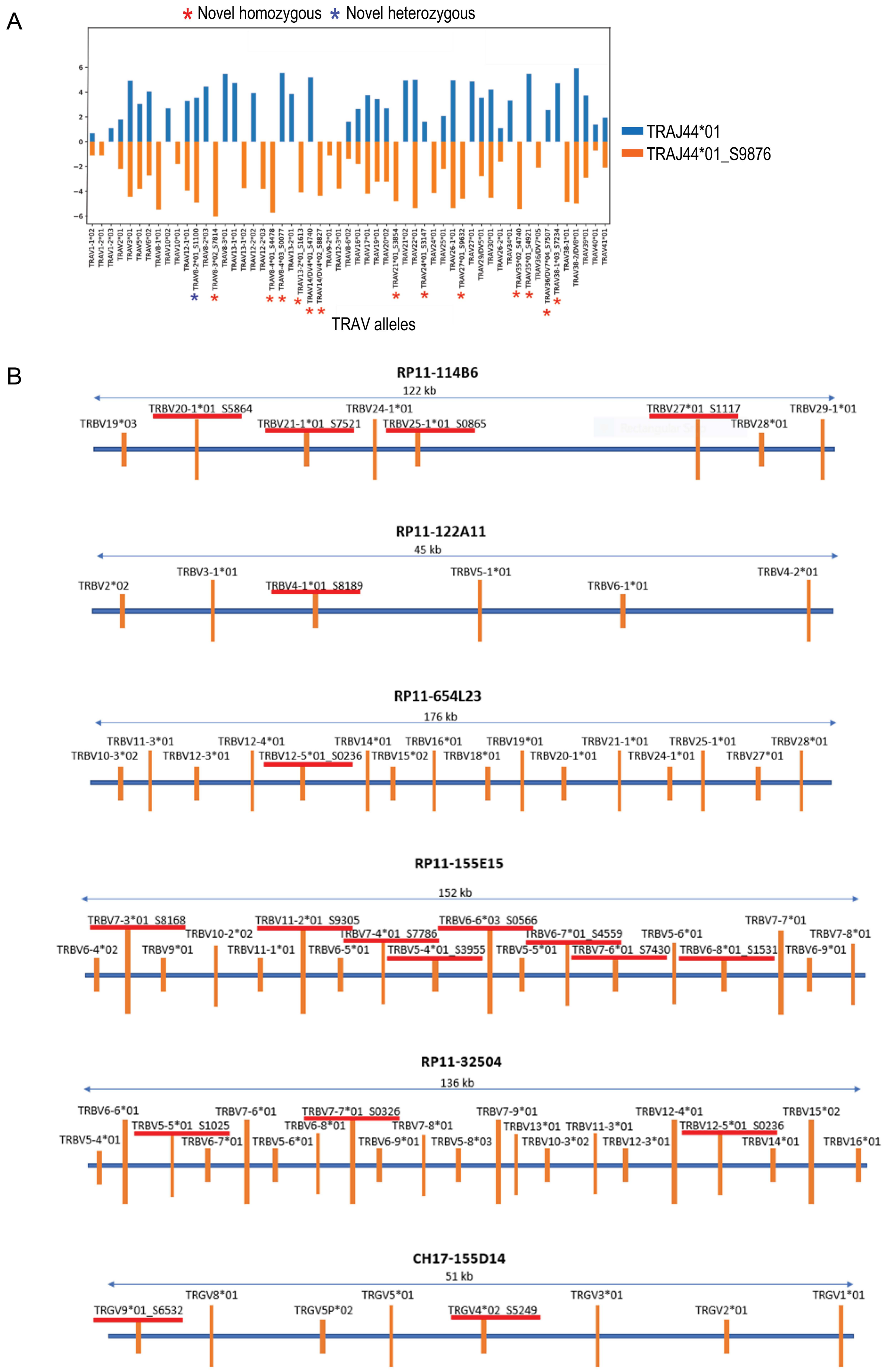

A

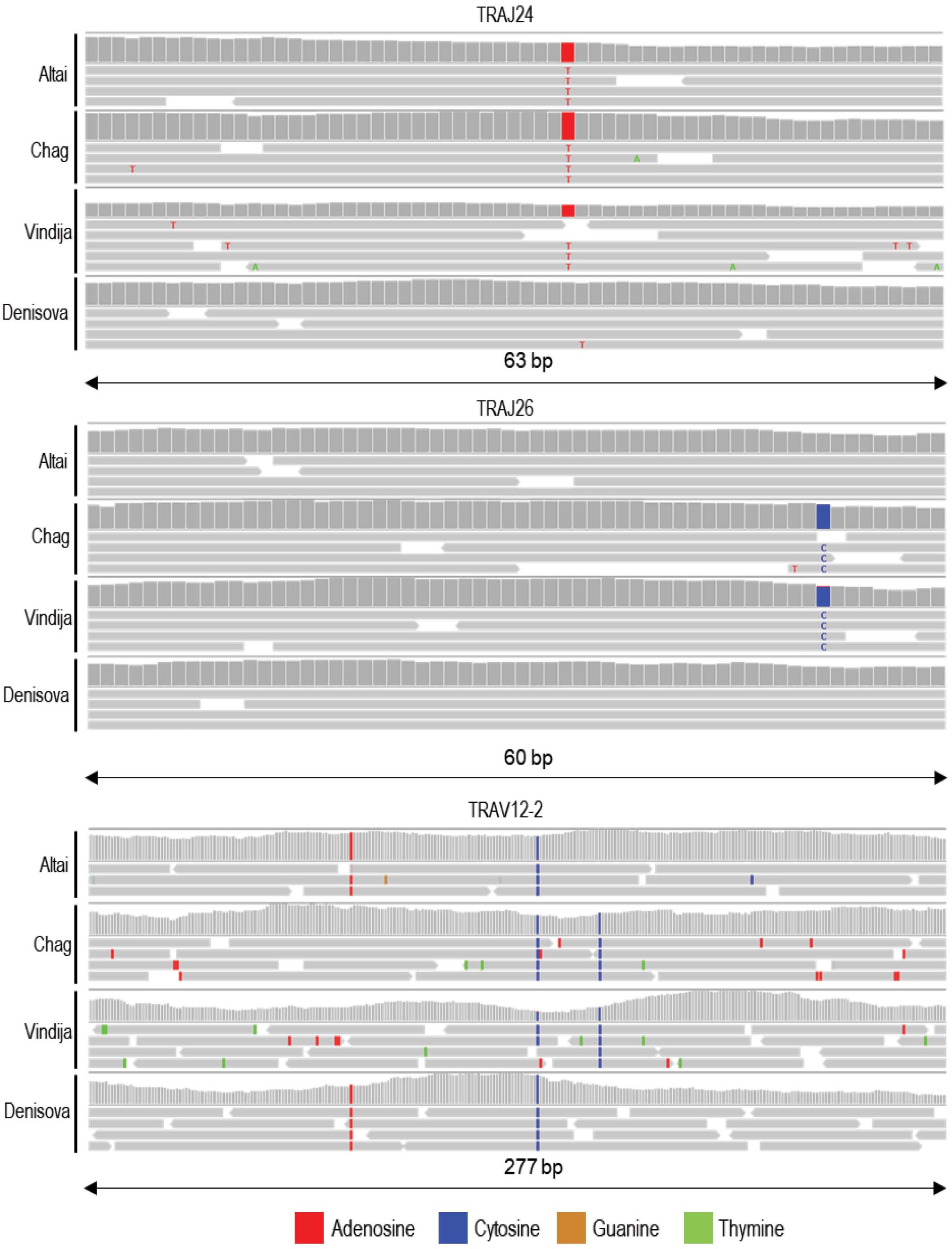


B

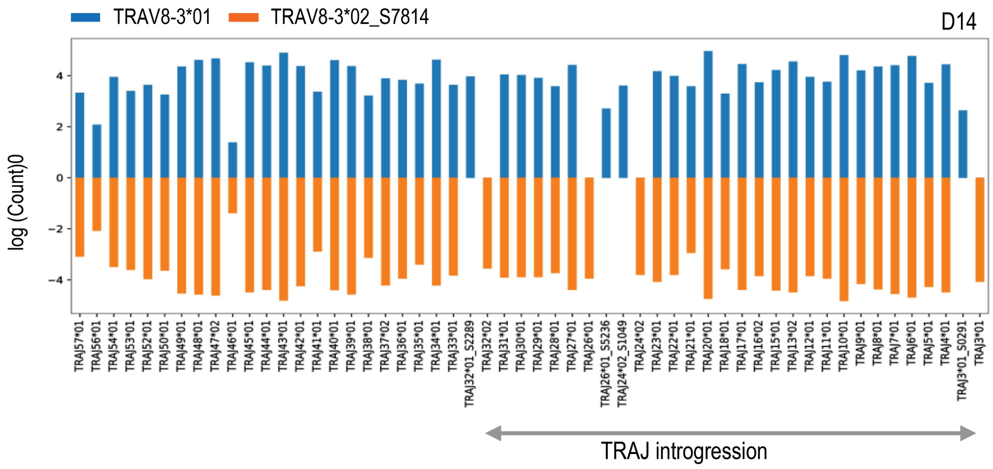


SI Figure 6

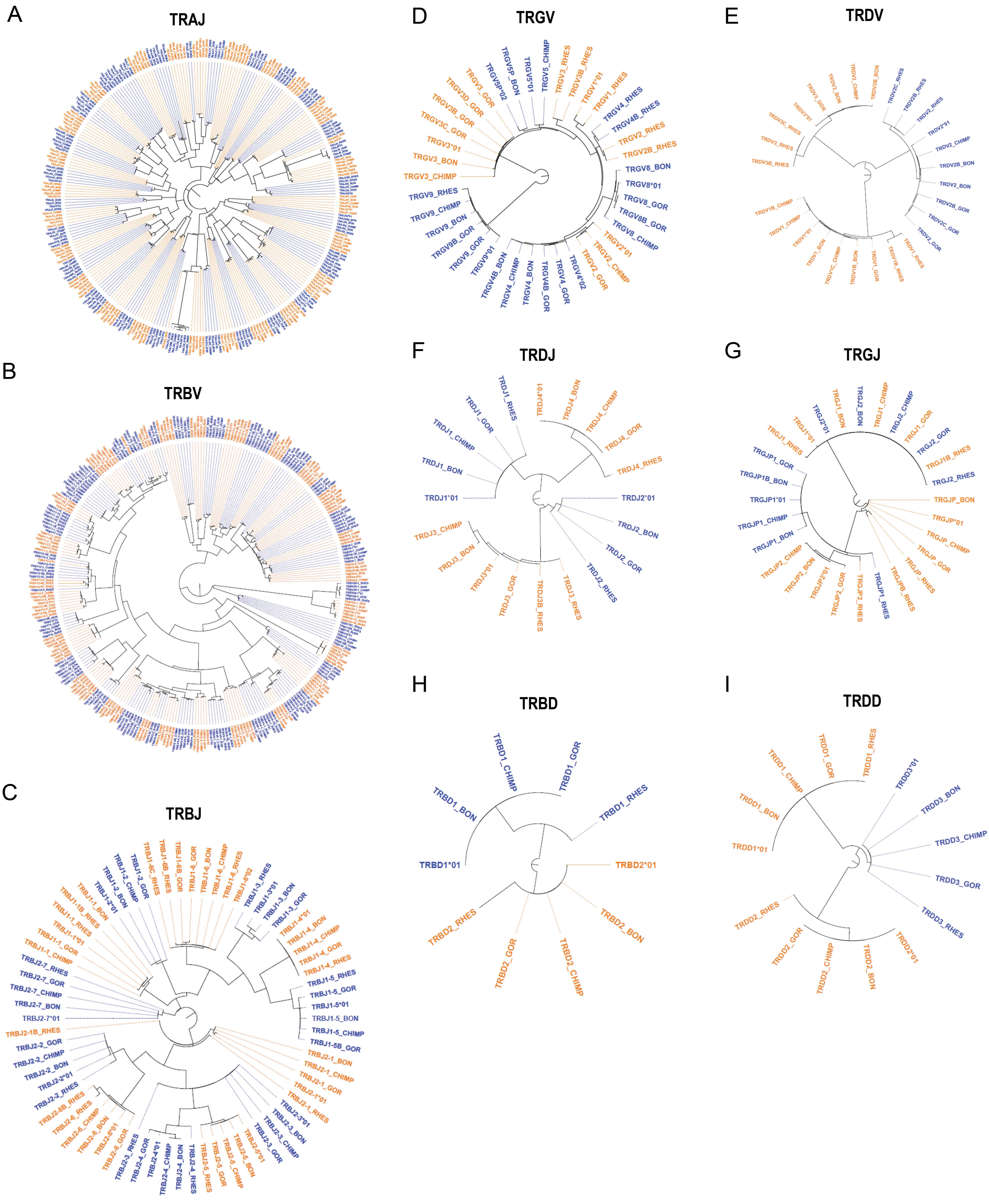
